# Talin controls the spatial distribution of vinculin tension in focal adhesions

**DOI:** 10.64898/2026.05.07.722930

**Authors:** Till Kallem, Yanyu Guo, Meghan K Reynolds, Neil J Ball, Muktesh Athale, Karen B Baker, Vasyl V. Mykuliak, Frans Ek, Silvia Aldaz-Casanova, Paula Turkki, Vesa P. Hytönen, Nicholas H Brown, Brenton D Hoffman, Jie Yan, Benjamin T Goult

## Abstract

Cells transmit force between the extracellular matrix and the actin cytoskeleton through integrin adhesion complexes centred on talin and vinculin. Vinculin binds talin through α-helical vinculin-binding sites (VBS) that are exposed when talin rod domains unfold under force. Dissecting the significance of this interaction has relied heavily on the A50I mutation in vinculin, which has been widely used as a talin-binding-null mutant. Here we show that although the A50I mutation abolishes binding to the α-catenin VBS, it retains nanomolar-affinity binding to multiple talin VBS. We therefore designed an improved mutant, I12K/A50I, that eliminates this residual talin binding. Biochemical assays and single-molecule stretching experiments demonstrate that I12K/A50I VD1 fails to bind talin even when VBS are exposed by force. Using vinculin tension and conformation sensors, we show that talin binding is required for efficient recruitment of vinculin to focal adhesions and for establishing spatial gradients of vinculin tension. However, vinculin can still experience mechanical load in the absence of talin binding. These results demonstrate that A50I is not a talin-binding-null and reveal that, while talin is not required for vinculin loading, it is essential for organising the spatial distribution of mechanical load within adhesion complexes.

## Introduction

Cells adhere to the extracellular matrix (ECM) through dynamic multi-protein complexes, termed integrin adhesion complexes (IACs) (Legerstee and Houtsmuller 2021; Petit and Thiery 2000; Bachmann et al. 2019). These complexes link the integrin ECM receptors to the actin cytoskeleton and are involved in mechanotransduction (Kuo 2013; Hoffman et al. 2011; Goult et al. 2018), signalling (Legerstee and Houtsmuller 2021; Goult et al. 2018) and cellular motility (Legerstee and Houtsmuller 2021; Petit and Thiery 2000; Kuo 2013).

Vinculin is a highly conserved protein with roles in both focal adhesions (FAs) and cell-cell contacts. At FAs, vinculin binds to talin, the core mechanotransducive protein in IACs which links the ECM-bound integrins to F-actin. Talin contains an N-terminal FERM domain that binds integrins and a large rod domain composed of 13 helical bundles (R1–R13) (Goult et al. 2013) that act as mechanical switches by unfolding under varying forces (Yao et al. 2016). These unfolding events reveal cryptic vinculin binding sites (VBS) (Fillingham et al. 2005; Yao et al. 2016; Yogesha et al. 2012; Del Rio et al. 2009), of which there exist at least 11 distributed throughout 9 of the 13 rod domains (Gingras et al. 2005) (**Fig. 1A**). Vinculin links exposed VBS in unfolded talin (and α-catenin at cell-cell junctions) (Choi et al. 2012; Peng et al. 2012) to F-actin (Humphries et al. 2007; Menkel et al. 1994; Carisey et al. 2013), reinforcing the talin tether between the ECM and cytoskeleton under force (Ciobanasu et al. 2014; Hirata et al. 2014; Giannone et al. 2003). Vinculin is primarily localised to FAs through its interaction with talin (Hirata et al. 2014; Zhang et al. 2008; Giannone et al. 2003; Wang et al. 2011; Goldmann et al. 1998; Pasapera et al. 2010) and potentially other binding partners (Bays and DeMali 2017).

**Figure 1.**
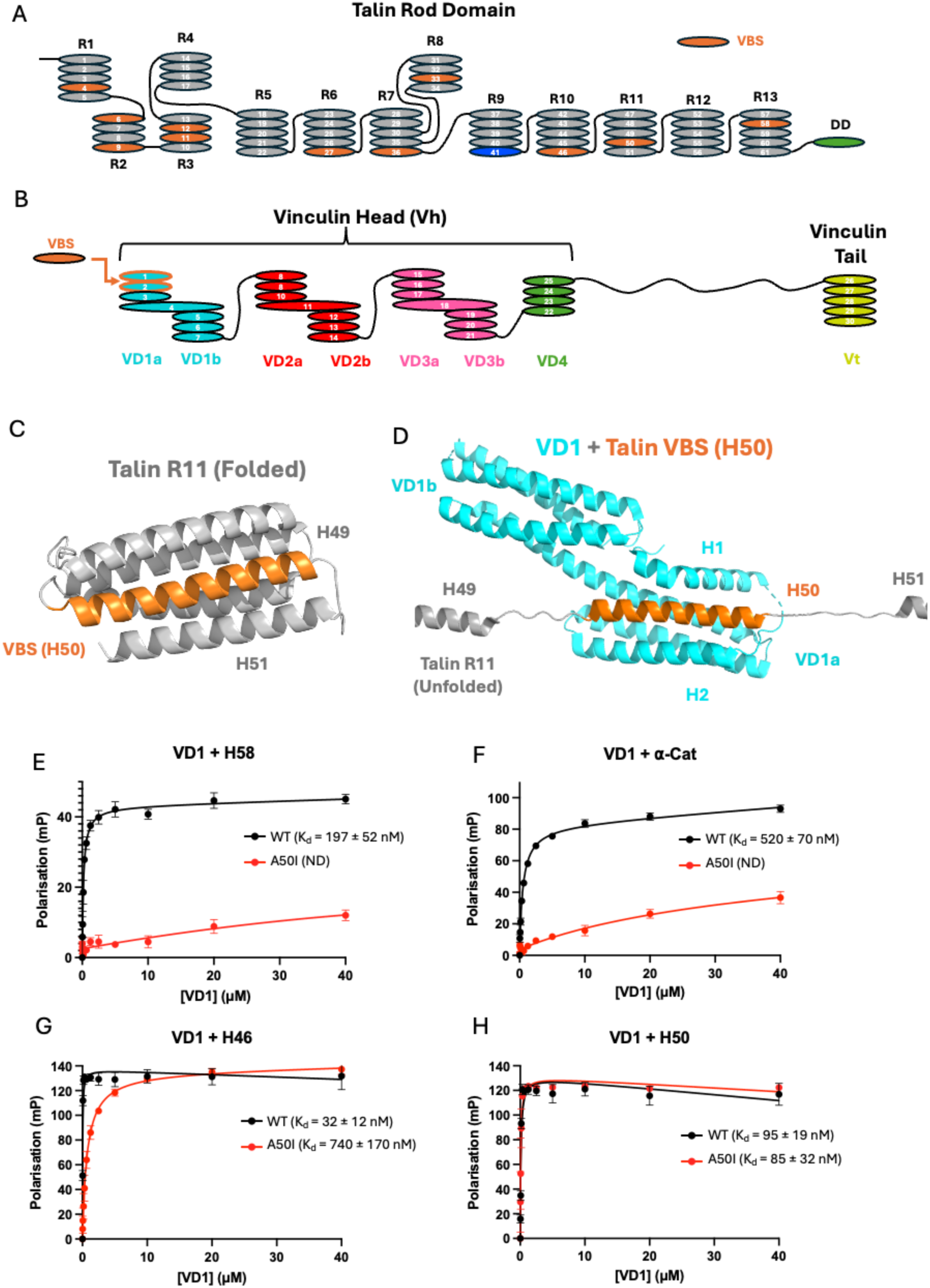
The A50I vinculin mutant retains binding to talin. **(A)** Schematic of the talin rod region showing the 13 rod domains (R1-R13). Vinculin-binding site (VBS) helices are highlighted in orange, the A-kinase anchoring (AKAP) helix in blue, and the dimerisation domain (DD) helix is shown in green. **(B)** Vinculin contains a single talin/α-catenin binding site in the VD1a domain (cyan). **(C-D)** Structures of talin-1 VBS helix 50 (H50; orange) from R11 folded within the R11 bundle **(C)** and a schematic model of mechanically exposed H50 bound to VD1 **(D)**. **(E-H)** Fluorescence polarisation assays of WT and A50I VD1 binding to talin and ⍺-catenin VBS peptides. **(E)** H58 binds tightly to WT but not to A50I VD1. **(F)** A50I prevents binding to the ⍺-catenin VBS. **(G)** A50I reduces the affinity of H46 by approximately 20-fold. **(H)** Binding to H50 is largely unaffected by the A50I mutation.

Vinculin is a 117 kDa protein which contains an N-terminal head domain (Vh) and a C-terminal tail domain (Vt) connected by a flexible linker (neck) (**Fig. 1B**). In mammals, Vh contains three double four-helix domains (VD1-3) and a single four-helix domain (VD4). Vt contains a single five-helix domain (Bakolitsa et al. 2004; Ziegler et al. 2006; Borgon et al. 2004). The ability of vinculin to crosslink is regulated by an auto-inhibitory interaction between Vh and Vt. In the inactive closed conformation, VD1 and VD4 interact strongly with Vt, leading to a compact structure that prevents many protein interactions (Bakolitsa et al. 2004; Borgon et al. 2004). Release of the head-tail interaction enables vinculin linker function in FAs and cell-cell contacts. In the active conformation, vinculin binds to talin and α-catenin through the VD1a domain (Izard et al. 2004; Papagrigoriou et al. 2004) while Vt binds F-actin (Johnson and Craig 1995; Menkel et al. 1994; Janssen et al. 2006), enabling the vinculin-mediated crosslink between talin and F-actin. Similarly to talin, the helical bundles of Vh also unfold under physiological forces, suggesting they act as mechanically induced binding sites that further buffer tension in adhesion complexes (Liu et al. 2025).

The binding of vinculin to VBS occurs through hydrophobic interactions between VD1a and the VBS helix (Papagrigoriou et al. 2004; Izard et al. 2004). Insertion of the VBS helix into the VD1a helical bundle is mediated by the separation of VD1a helices 1 and 2. The resultant complex is a 5-helix bundle with a hydrophobic core that is reminiscent of a talin rod domain prior to mechanical unfolding (**Fig. 1C-D**) (Bakolitsa et al. 2004; Izard et al. 2004; Papagrigoriou et al. 2004; Bass et al. 2002; Mykuliak et al. 2024; Gingras et al. 2005; Yogesha et al. 2012).

Mutations in vinculin have been developed to disrupt the vinculin-mediated mechanical linkage between talin and actin. A point mutation in Vt (I997A) uncouples vinculin from F-actin (Thompson et al. 2014). Similarly, an A50I point mutation in the VD1a domain disrupts binding to vinculin-binding sites (VBS) by stabilising the closed conformation of the VD1a binding groove (Bakolitsa et al. 2004; Cohen et al. 2006). This mutant has therefore been widely used as a talin-binding-null version of vinculin. However, the efficacy of A50I in preventing binding to talin remains unclear in the literature. Broadly, these discrepancies largely occur between *in vitro* and *in cellulo* experiments. Biochemical studies, typically conducted in solution and without loading applied to talin, routinely show a lack of interaction between talin and A50I vinculin or A50I VD1 (Humphries et al. 2007; Cohen et al. 2006; Hui Chen et al. 2006; Peng et al. 2012). Vinculin (-/-) fibroblasts reconstituted with A50I vinculin show reduced force-generation, perturbed FA maturation, and inefficient directed migration, suggesting strong perturbation to cell function (Baumann et al. 2023). However, FRET-based biosensor studies in cells have shown that A50I vinculin can still undergo activation and maintain, or even increase, mechanical loading (Hui Chen et al. 2005; Grashoff et al. 2010). Together, these observations raise the possibility that A50I reduces, but does not abolish, force-sensitive vinculin–talin interactions in cells.

Despite its widespread use as a talin-binding-null construct, it remains unclear whether the A50I mutation fully abolishes vinculin binding to talin. Here we show that A50I vinculin retains binding to multiple talin VBS and therefore is not a talin-binding null. We introduce an improved mutant, I12K/A50I, that eliminates this residual talin binding. Using this tool, we demonstrate that while vinculin can still be mechanically loaded in focal adhesions in the absence of talin binding, the talin–vinculin interaction is critical for efficient recruitment of vinculin and for establishing spatial gradients of tension across adhesions. Together, these findings reveal that talin spatially organises vinculin tension and force transmission within integrin adhesion complexes.

## Results

### A genetic screen identifies mutations that disrupt VD1 activity

Whereas overexpression of full-length vinculin does not induce phenotypes in *Drosophila*, the expression of constitutively active truncated forms, such as Vh or VD1, causes lethality (Maartens et al. 2016). To investigate the basis of this lethality, we performed a genetic screen to identify mutations in VD1 that suppress lethality (see Materials and Methods). Screening approximately 66,000 mutagenised flies yielded four missense mutations in conserved VD1 residues that eliminated lethality: P15L, A50V, G58E and G118E. Notably, one mutation altered A50 to valine, a residue similar to the A50I mutant designed to stabilise the unbound conformation of VD1 (Bakolitsa et al. 2004). Mutating A50 to isoleucine was not possible with the mutagen used. Unexpectedly, despite suppressing lethality, these mutations did not impair VD1 recruitment to integrin adhesion sites in embryonic muscles or to cell-cell junctions in imaginal discs (data not shown). This indicates that lethality is not caused by altered localisation of VD1, but instead reflects changes in its interactions with ligands such as talin. The retention of focal adhesion recruitment by A50V VD1 suggested that this mutation, and by extension A50I, may not block binding to all VBS.

### A50I VD1 still binds to talin

To test whether the A50I mutation prevents binding of vinculin to individual VBS, we used fluorescence polarisation (FP) to measure binding of WT and A50I VD1 to fluorescein-tagged synthetic peptides corresponding to mouse talin-1 VBS helices (H46 from R10, H50 from R11, and H58 from R13; **Fig. 1A**) and the mouse ⍺**-**catenin VBS (**Fig. 1E-H**). As expected, all four VBS peptides bound to WT VD1 with nanomolar affinity. The A50I mutation abolished binding to H58 (ND, **Fig. 1E**) and strongly reduced binding to the ⍺-catenin VBS (ND, **Fig. 1F**). Binding to H46 was reduced ∼20-fold but remained sub-micromolar (K_d_ = 740 ± 170 nM, **Fig. 1G**). Unexpectedly, binding to H50 was largely unchanged (A50I: K_d_ = 85 ± 32 nM versus WT: 95 ± 19 nM, **Fig. 1H**). These results show that although the A50I mutation disrupts VD1 binding to ⍺**-**catenin, it only partially disrupts binding to talin VBS. Thus, A50I VD1 behaves as an α-catenin-binding null but not a talin-binding null.

### The structure of H50 in complex with A50I VD1

Because binding of H50 was unaffected by the A50I mutation, we sought to understand how this interaction is accommodated structurally. The A50I VD1-H50 complex crystallised readily, and the crystals diffracted to 1.5 Å resolution. The structure was solved by molecular replacement using the WT VD1-H50 complex (PDB ID: 4DJ9, Yogesha et al. 2012) as the search model. A single complex was present in the asymmetric unit in space group P2_1_2_1_2_1_.

Helices 1 and 2 of A50I VD1 are separated relative to each other, facilitating insertion of the H50 helix (**Fig. 2**). Structural alignment of the A50I VD1-H50 structure with WT VD1-H50 (4DJ9; (Yogesha et al. 2012; Gingras et al. 2005) revealed remarkable similarity (RMSD = 1.55 Å across 1865 atoms) (**Fig. 2A-B**). These results show that the A50I mutation does not alter the bound conformation of the VD1-H50 complex.

**Figure 2.**
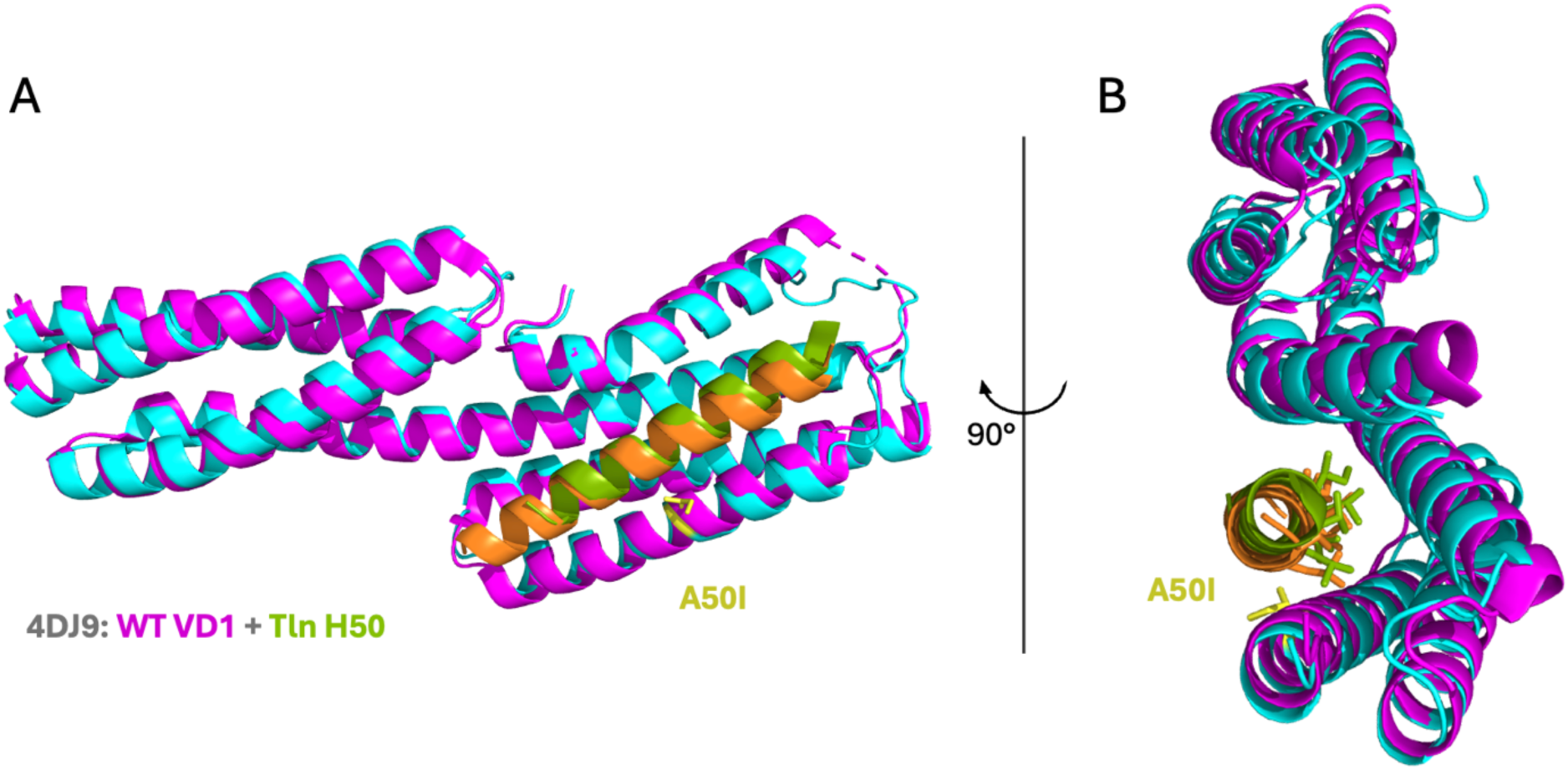
Crystal structure of A50I VD1 bound to talin-1 H50. **(A-B)** Structural alignment of the A50I VD1-H50 crystal structure (cyan/orange) with the WT VD1-H50 (PDB ID: 4DJ9; magenta/green (Yogesha et al. 2012)) shown in two orientations.

The preserved structure of the A50I VD1–H50 complex suggested that features of the VBS helix may permit binding despite the A50I mutation. Early exploratory molecular dynamics simulations of VBS–vinculin interactions suggested a role for N-terminal hydrophobic residues in the initial engagement with VD1 prior to VBS insertion. Guided by this, comparison of talin VBS sequences suggested that the hydrophobic character of the VBS helix may contribute to this behaviour. Among the 11 talin VBS, H50 contains the most hydrophobic N-terminus (VVLI) (**Fig. 3A**). This hydrophobic segment is well positioned to dock into the hydrophobic groove of unbound VD1, including the region surrounding A50I (**Fig. 3C–D**), potentially lowering the energetic barrier to VBS insertion (Mykuliak et al. 2024). This may explain why the binding affinity of H50 is largely unaffected by the A50I mutation (**Fig. 3B**).

**Figure 3.**
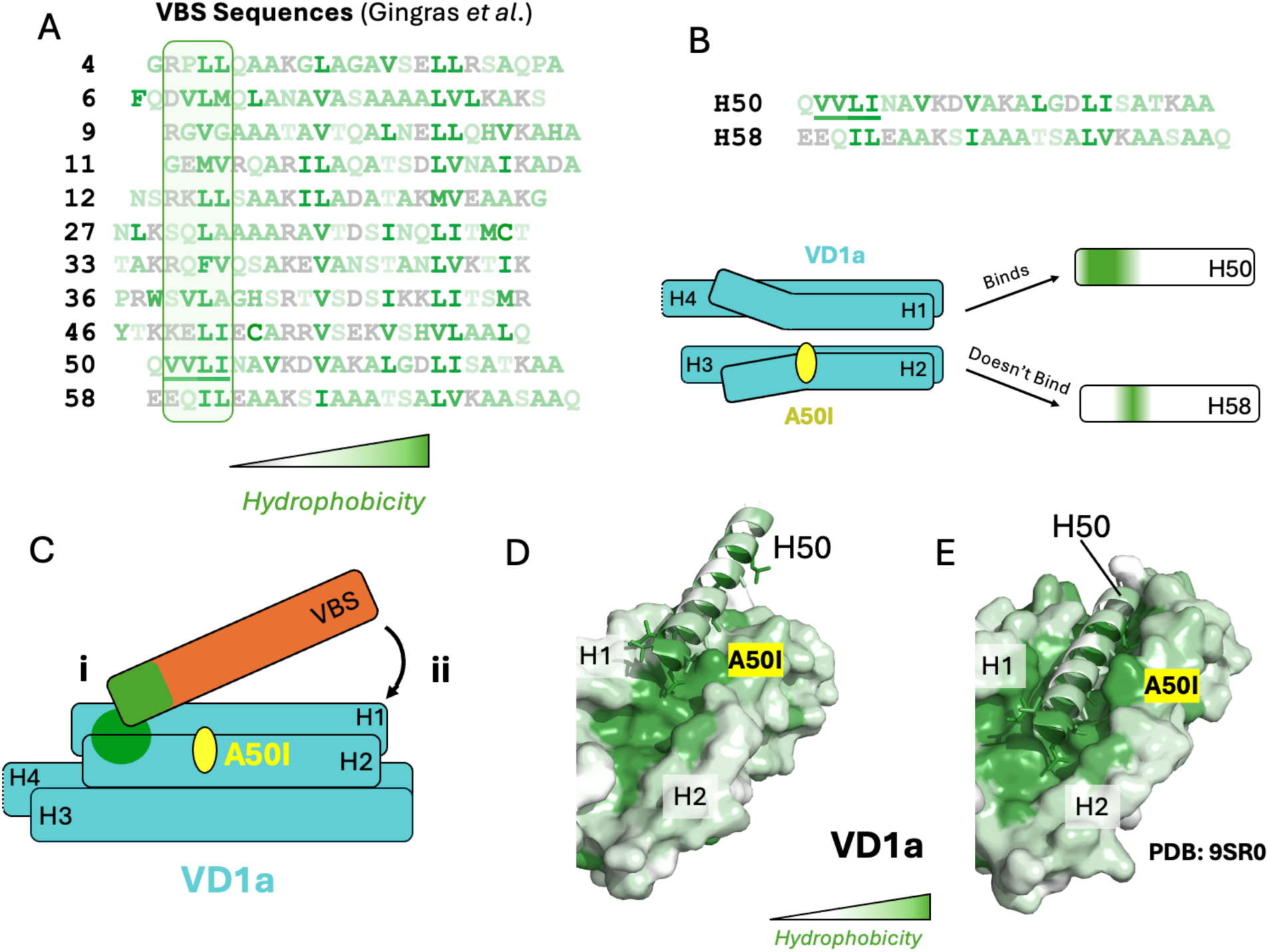
Hydrophobic N-termini enable talin VBS binding to A50I VD1. **(A)** Talin-1 VBS sequence alignment from (Gingras et al. 2005) coloured by hydrophobicity using the AAindex scale (entry FASG890101; (Nakai et al. 1988)). The highly hydrophobic N-terminal four residues of H50 (VVLI) are underlined and compared with other VBS. **(B)** Cartoon model illustrating that VBS helices with more hydrophobic N-termini (e.g., H50) can bind A50I VD1. **(C)** Cartoon model of (i) VBS N-terminus docking into VD1a binding groove along hydrophobic patch before (ii) inserting lengthwise into binding groove. **(D)** Surface representations of H50 docking into the VD1a hydrophobic patch. The N-terminus of H50 interacts with the hydrophobic patch adjacent to the A50I site on VD1a. **(E)** Surface representation of the A50I VD1-H50 final complex (PDB ID: 9SR0) coloured by hydrophobicity as above.

### Improving on A50I to generate a talin-binding-null vinculin

Having shown that the A50I mutation is ineffective at preventing talin binding, we used A50I as a foundation to design a mutant that abolishes detectable talin binding. Comparison of available talin-vinculin crystal structures revealed that isoleucine 12 lies adjacent to a conserved positively charged residue present in most VBS when bound (Gingras et al. 2005) (**Fig. S1A**). This residue forms ion-dipole interactions with nearby serine and threonine residues on VD1, as observed in crystal structures of VD1 in complex with H11 (PDB ID: 1ZVZ, Gingras et al. 2005), H36 (PDB ID: 1ZW3, Gingras et al. 2005), H50 (PDB ID: 4DJ9, Yogesha et al. 2012) and H58 (PDB ID: 1ZW2, Gingras et al. 2005) (**Fig. S1B**).

We therefore introduced a positive charge (I12K) into A50I VD1 with two intended effects: (i) disruption of hydrophobic interactions that stabilise the VBS-VD1 complex, and (ii) electrostatic repulsion between the introduced lysine and the conserved positively charged residue in the VBS helix (**Fig. 4A-B**). We reasoned that, in combination, A50I would stabilise the unbound state while I12K would destabilise the bound state. Together, these mutations are expected to prevent binding of the remaining talin VBS that still interact with A50I VD1.

**Figure 4.**
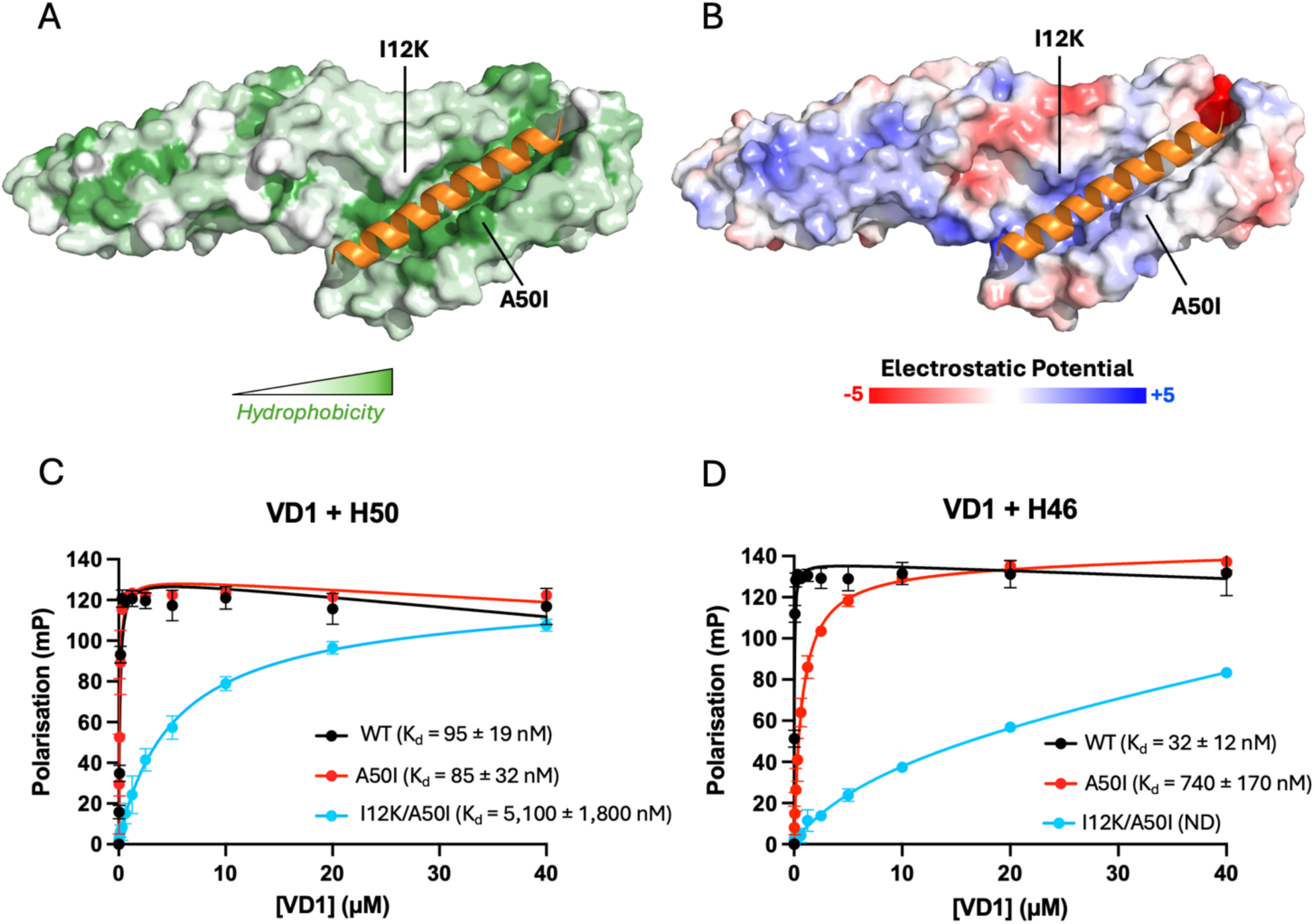
Rational design of a talin-binding-null vinculin mutant. **(A–B)** Surface representations of the A50I VD1–H50 crystal structure with the I12K mutation modelled in silico, coloured by **(A)** hydrophobicity using the AAindex database (entry FASG890101; (Nakai et al. 1988)) in PyMOL and **(B)** electrostatic surface potential, showing increased positive charge within the VBS-binding groove. H50 is shown in orange. **(C-D)** Fluorescence polarisation assays showing binding of WT, A50I, and I12K/A50I VD1 to **(C)** H50 and **(D)** H46.

Circular Dichroism (CD) spectra of VD1 containing the A50I and I12K mutations, individually or in combination, were indistinguishable from WT VD1, indicating that these mutations do not disrupt protein folding (**Fig. S2A**). We next assessed thermal stability by monitoring ellipticity at 222 nm while increasing the temperature from 20-90°C. WT VD1 exhibited biphasic melting (**Fig. S2B**) with distinct transitions corresponding to VD1a (T_m_ = 44.8°C) and VD1b (T_m_ = 65.1°C) (**Fig. S2C**). These biphasic profiles were preserved in all mutants. A50I stabilised VD1a (T_m_ = 54°C), whereas I12K destabilised it (T_m_ = 40°C). In the double mutant, these effects were balanced, with VD1a melting at approximately the same temperature as WT (T_m_ = 44°C).

Addition of H46 or H50 stabilised the lower-temperature transition of WT VD1, confirming its assignment to VD1a (**Fig. S2D**). In contrast, only H50 stabilised A50I VD1a (**Fig. S2E**), consistent with reduced H46 binding. The I12K mutation alone had minimal effects (**Fig. S2F**), whereas the I12K/A50I double mutant showed no detectable stabilisation with either H46 or H50 (**Fig. S2G**).

FP experiments confirmed that the I12K/A50I double mutant substantially weakened binding to H50 (∼60-fold reduction; K_d_ = 5.1 ± 1.8 μM; **Fig. 4C**) and abolished binding to H46 (ND; **Fig. 4D**). The combined mutations acted synergistically, as I12K alone produced only modest reductions in affinity for H50 (K_d_ = 127 ± 39 nM; **Fig. S3A**) and H46 (K_d_ = 800 ± 12 nM; **Fig. S3B**). Consistent with these findings, GST pulldown assays showed no detectable interaction (**Fig. S4**).

### Single molecule analysis demonstrates that I12K/A50I VD1 does not bind talin under tension

In a previous study, we used magnetic tweezers to characterise VD1 binding to the full talin-1 rod (R1-R13) and showed that 9 of the 13 talin rod domains bind VD1 when mechanically unfolded (Yao et al. 2016). VD1 binding to exposed VBS prevents domain refolding, effectively locking talin domains in the unfolded state (Yao et al. 2014). To test whether A50I and I12K/A50I VD1 bind VBS exposed under force, we applied this single-molecule stretching assay to R1–R13 in the presence of WT, A50I or I12K/A50I VD1 (**Fig. 5A**).

**Figure 5.**
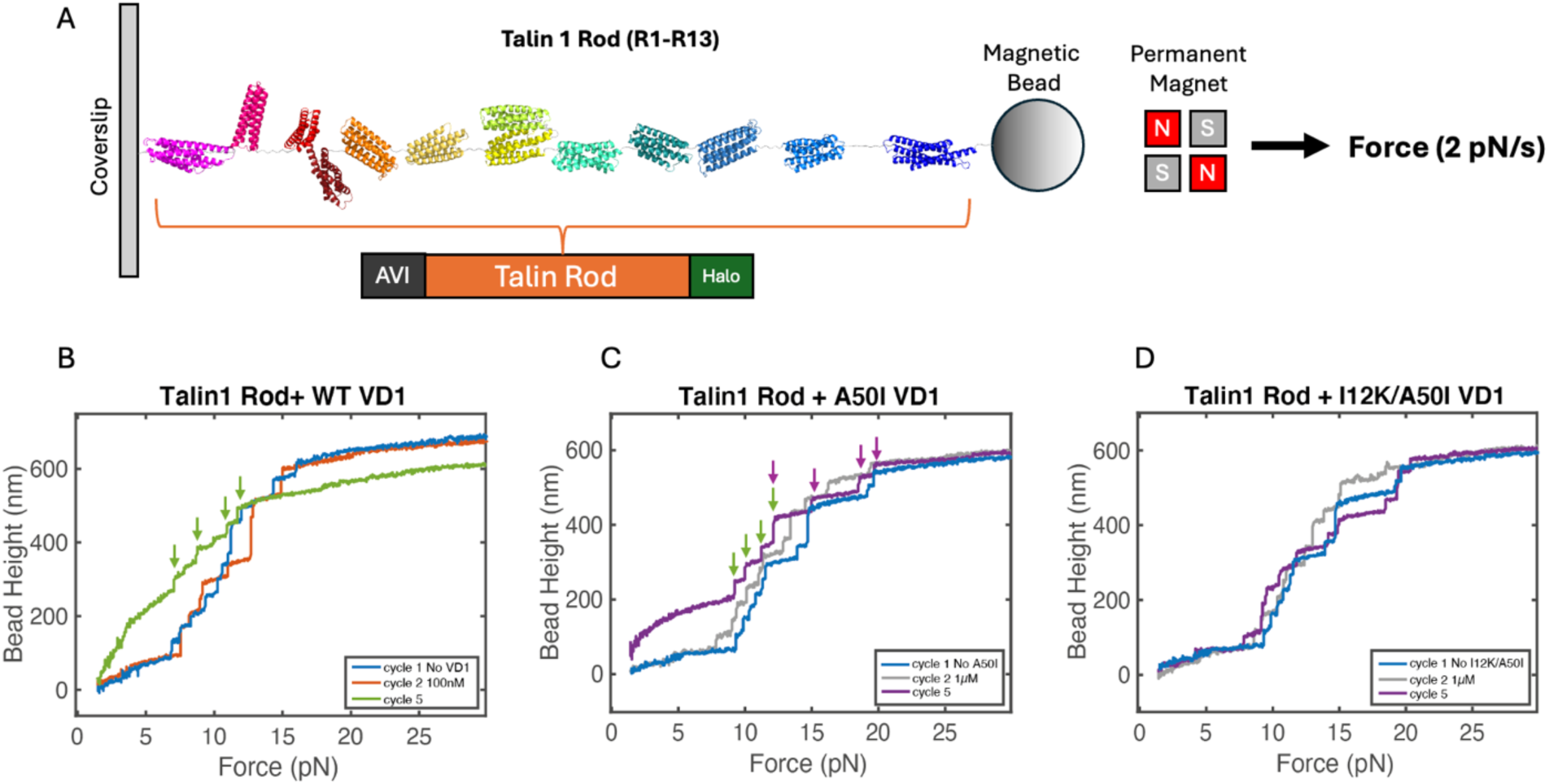
Single-molecule stretching of the talin rod in the presence of VD1. **(A)** Experimental setup. The mouse talin-1 rod (R1-R13), containing an N-terminal Avi-tag and a C-terminal Halo-tag, was tethered between a glass coverslip and a 2.8 µm paramagnetic bead using Halo-tag and Avi-tag/streptavidin chemistry. **(B-D)** Representative unfolding force-extension curves recorded at a loading rate of 2 pN s^-1^ in the presence of **(B)** WT VD1 (100 nM), **(C)** A50I VD1 (1 µM), and **(D)** I12K/A50I VD1 (1 µM). The four unfolding events that persist in the presence of WT VD1 are indicated by green arrows. Additional unfolding events observed in the presence of A50I VD1 are indicated by purple arrows. In contrast, unfolding curves recorded in the presence and absence of I12K/A50I VD1 were indistinguishable, consistent with loss of detectable talin binding.

During the first stretching event, 12 unfolding events were observed, corresponding to the 13 talin rod domains, with R7 and R8 unfolding together as a single event at 12-15 pN (Yao et al. 2016) (**Fig. 5B**). Following relaxation to 1 pN for 100 s to allow refolding, subsequent stretching cycles were used to assess VD1 binding. In the presence of WT VD1 (100 nM), the number of unfolding events decreased from 12 to 4, as VD1 binding to the nine VBS-containing rod domains prevented their refolding (**Fig. 5B**) (Yao et al. 2016). The remaining four domains (R4, R5, R9, and R12) lack VBS (Gingras et al. 2005).

In contrast, A50I VD1 showed markedly reduced binding. At 100 nM, a single domain was prevented from refolding (**Fig. S5B**), consistent with a high-affinity interaction, likely with H50 in R11. At higher concentrations (1 μM), up to five talin rod domains were prevented from refolding in each cycle (**Fig. 5C**), indicating that additional VBS can bind A50I VD1, albeit with lower affinity or shorter lifetimes. Based on the previous binding assays, these A50I-VD1-binding domains likely include R11 (H50) and R10 (H46) (**Fig. 1G-H**).

Strikingly, I12K/A50I VD1 failed to inhibit talin refolding even at high concentrations. At 1 μM, all talin rod domains refolded during the 100 s holding period at 1 pN in most cycles (Fig. 5D, Fig. S5D), indicating that I12K/A50I is unable to bind mechanically exposed VBS under force. Together, these experiments show that, whereas A50I VD1 retains force-dependent binding to multiple talin VBS, the I12K/A50I mutant abolishes vinculin binding even when VBS are mechanically exposed.

### FRET-based tension sensor analysis reveals that talin-binding-null vinculin remains under tension in FAs

To determine how disruption of talin binding affects vinculin function, we used a suite of vinculin Förster resonance energy transfer (FRET)-based biosensors expressed in vinculin (-/-) mouse embryonic fibroblasts (MEFs). Specifically, vinculin conformation sensors (CS) were used to assess vinculin activation state, as has been done previously when characterising vinculin mutations (Shoyer et al. 2023; Chirasani et al. 2023). In these sensors, release of vinculin head-tail inhibition leads to an active, open conformation of vinculin and reduction in the sensors’ FRET efficiency (**Fig. S7A-B**) (Hui Chen et al. 2005). Likewise, vinculin tension sensors (TS) were used to assess vinculin loading, with applied tension extending the sensors’ tension-sensing module and leading to lower FRET efficiencies (**Fig. 6A-B**) (Grashoff et al. 2010). Both conformation and tension sensors can also be used to analyse vinculin localisation through their acceptor intensity, which is insensitive to FRET. In this study, the sensor suite included WT, A50I, and novel I12K-A50I vinculin CS and TS (Grashoff et al. 2010; Rothenberg et al. 2018). The I997A mutation, which disrupts vinculin-actin interaction (Thompson et al. 2014), was also introduced into the I12K/A50I vinculin sensors to serve as a zero-tension-control.

**Figure 6.**
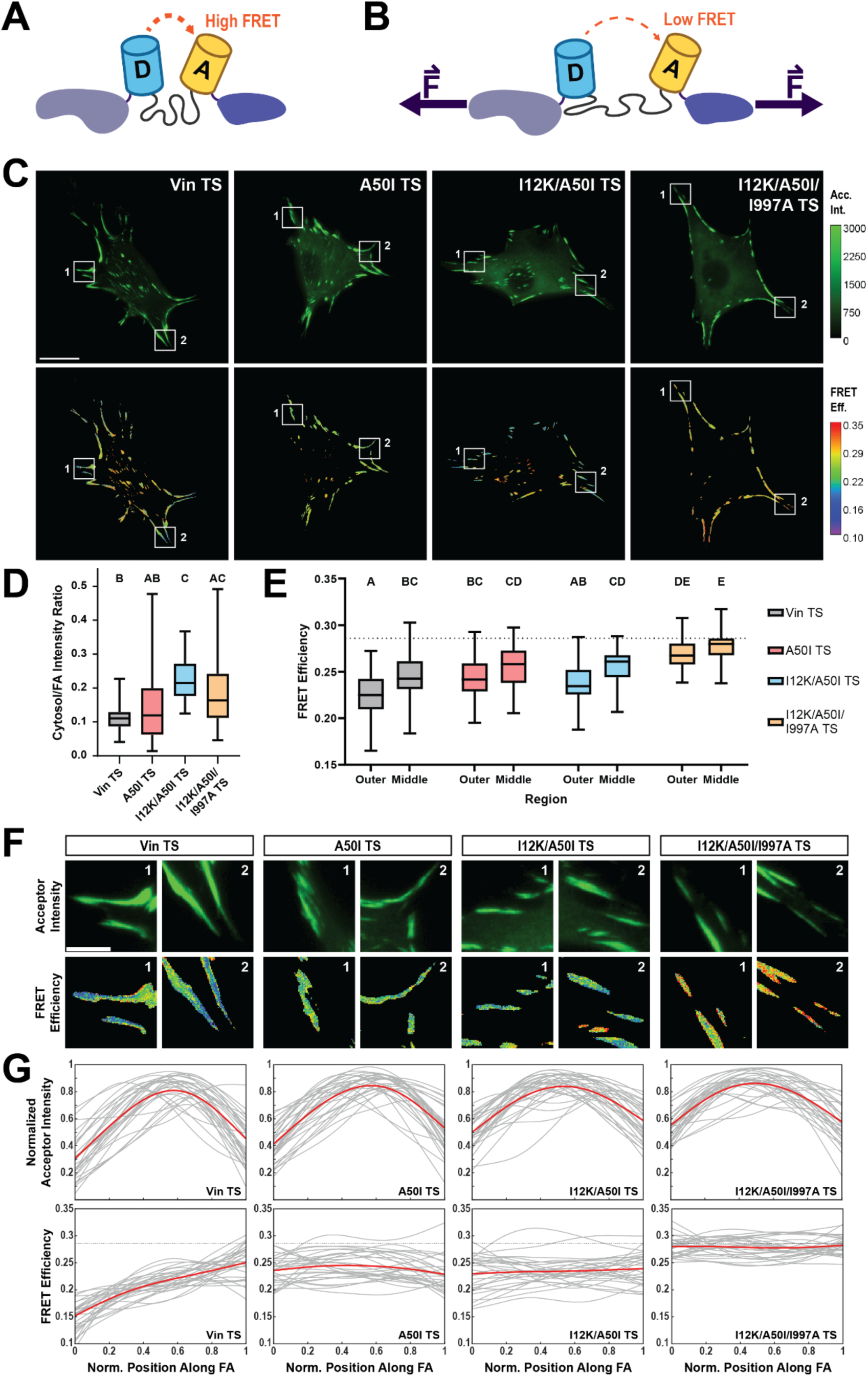
Talin binding promotes vinculin recruitment and spatial organisation of tension in focal adhesions. **(A-B)** Schematic diagram of the vinculin tension sensor under (A) low/no tension (high FRET) and (B) high tension (low FRET). **(C)** Representative images of acceptor intensity (top) and masked FRET efficiency (bottom) in vinculin (-/-) MEFs expressing vinculin tension sensor constructs. Scale bar, 20 µm. **(D-E)** Box and whisker plots showing cell-averaged cytosol-to-FA acceptor intensity ratios **(D)** and cell-averaged FRET efficiency in outer and middle regions of the cell **(E)**. Letters above box plots represent statistical significance, with box plots with no letter in common being significantly different from one another at p < 0.05. Statistical analysis was performed using one-way ANOVA followed by Steel-Dwass or Tukey Kramer post hoc tests, respectively. The dashed line indicates the previously established unloaded FRET efficiency. Data represent n = 59, 32, 29, and 36 biologically independent cells, collected across N = 3 independent experimental days. **(F)** Zoomed regions of FAs in the outer region of the cell corresponding to the boxed regions in (C). Scale bar, 5 µm. **(G)** Line scans of acceptor intensity (top) and FRET efficiency (bottom) across individual FAs from distal (position 0) to proximal (position 1) tips. Grey lines represent scans from 29 FAs across three independent experimental days, red lines indicate the averaged profile, and dotted lines indicate the previously established unloaded FRET efficiency.

We began by evaluating the impact of talin-binding on vinculin localisation through acceptor intensity analysis. Initial qualitative assessment confirmed that all constructs could successfully localise to FAs (**Fig. S7A-C, Fig. 6C**). Consistent with this, residual recruitment of A50I VD1 but not A50I/I12K VD1 to FAs was also observed in cells even in the presence of endogenous vinculin (**Fig. S8**). However, talin-binding-deficient mutants appeared to have a larger fraction of vinculin present in the cytosol. To quantify these localisation differences, we compared the ratio of vinculin in the cytosol to vinculin at focal adhesions across constructs (**Fig. S5A**). This analysis revealed that ablation of residual talin-binding with I12K/A50I significantly increased the proportion of vinculin within the cytosol as compared to FAs for both sensor types (**Fig. S7D, Fig. 6D**). These data demonstrate that talin-binding is not strictly required, but plays a critical role in vinculin recruitment to FAs. This finding is consistent with previous studies implicating talin in vinculin localisation (Zhang et al. 2008), while isolating this effect without the broader disruption to adhesions caused by talin loss.

### FRET-based biosensor analysis reveals talin-vinculin interactions are essential for tension gradients in mature FAs

The sensor suite was then used to evaluate vinculin function at FAs. Notably, vinculin function can be regulated at both the cell and focal adhesion level (Grashoff et al. 2010; LaCroix et al. 2018; Blakely et al. 2014; Sarangi et al. 2017; Rothenberg et al. 2018; Aytekin et al. 2026). For instance, cell-level regulation of vinculin loading displays various relationships to traction force generation and plays important roles in cell polarisation and migration (Grashoff et al. 2010; Aytekin et al. 2026). To assess this cell-level regulation, we examined how vinculin function changes with FA distance from the cell centroid (**Fig. S6B**). This analysis specifically focused on FAs in the outer and middle regions of the cell, since FAs closest to the cell centroid contain unloaded vinculin (Rothenberg et al. 2018; Aytekin et al. 2026). Meanwhile, variations in vinculin load across individual FAs may play important roles in force transmission, with vinculin load and FA traction forces following similar spatial distributions (LaCroix et al. 2018; Blakely et al. 2014; Aytekin et al. 2026). This FA level regulation was also assessed through analysis of individual, peripheral FAs, as we and others have done previously (LaCroix et al. 2018; Sarangi et al. 2017).

Such evaluation with the vinculin CS suite showed that WT and all vinculin variants are open at FAs, relative to the inactive FRET efficiency value of 0.4 (**Fig. 6C**) (Chirasani et al. 2023; Shoyer et al. 2023). Cell-level analysis of these FRET efficiencies revealed an activation gradient in wildtype vinculin (**Fig. 6E**), but not in the variants. At the focal adhesion level, qualitative analysis illustrated that peripheral focal adhesions across all constructs had uniform FRET efficiency values and, therefore, degree of activation (**Fig. S7F**). Together, these results demonstrate that talin-binding is not required for vinculin activation at FAs, but promotes it at the cell edge.

The vinculin TS suite also revealed that WT and all vinculin variants, except the zero-tension-control construct, which reports the expected unloaded FRET efficiency of 28.6, bear tension at FAs (**Fig. 6C**). This finding matches previous results that showed vinculin A50I is still under tension, although at reduced levels compared to WT vinculin, supporting a talin-independent force transmission pathway (Rothenberg et al. 2018). Analysis at the cell-level revealed a gradient of tension in WT vinculin (**Fig. 6E**). Interestingly, this cell-level tension gradient was eliminated in A50I vinculin but restored upon elimination of residual talin-binding with I12K/A50I vinculin (**Fig. 6E**). This suggests that the weakened interactions between A50I vinculin and talin leads to a reduction in vinculin loading at the cell periphery. Meanwhile, at the individual FA-level, TS variants were similarly distributed throughout FAs, but clear differences in FRET distributions are apparent (**Fig. 6E-F**). Comparison revealed that the vinculin-talin interaction is required for vinculin tension gradients across individual FAs. Moreover, this analysis illustrates that talin-binding-deficient constructs cannot reach peak vinculin tension within the membrane proximal tips of mature FAs. Together, these findings suggest that, while the vinculin-talin interaction is not required for vinculin loading, it plays a key role in the tuning of vinculin loading within mature FAs near the cell periphery.

## Discussion

How vinculin becomes mechanically engaged within integrin adhesion complexes and adherens junctions remains a central question in mechanotransduction. Dissecting this process has relied heavily on point mutations designed to disrupt vinculin binding to talin and α-catenin. The alanine-to-isoleucine substitution at position 50 (A50I) has therefore been widely used as a talin-binding-null mutant of vinculin.

Prior to this study, the ability of α-catenin to bind A50I vinculin was unclear. Some studies reported that A50I abolished binding to α-catenin (Choi et al. 2012), whereas others suggested residual interaction or tension loading (Peng et al. 2012; Morales-Camilo et al. 2024). Our fluorescence polarisation measurements resolve this ambiguity, demonstrating that A50I VD1 does not bind the exposed α-catenin VBS.

In contrast, A50I only partially disrupts binding to talin VBS. Biochemical assays show that A50I VD1 retains nanomolar affinity for certain talin VBS helices, including H46 (R10) and H50 (R11). The crystal structure of the A50I VD1-H50 complex reveals an interaction that is nearly identical to that observed for WT VD1 bound to H50 (PDB 4DJ9, Yogesha et al. 2012). This is consistent with the proposed mechanism of the A50I mutation, which increases the activation energy required for VBS insertion by strengthening hydrophobic interactions between helices 1 and 2 of VD1a (Bakolitsa et al. 2004), but does not alter the final bound conformation once insertion occurs. Consequently, if a VBS helix is sufficiently hydrophobic to overcome this energetic barrier, binding can still occur.

These observations indicate that VBS binding to VD1 depends on both the energetic barrier for helix insertion and stabilisation of the bound complex. We therefore designed an improved mutant, I12K/A50I vinculin, that behaves as a talin-binding null. In this design, A50I increases the energy required for VBS insertion, whereas I12K introduces electrostatic repulsion with a conserved positively charged residue in the VBS helix. When combined, the two mutations act synergistically to prevent VBS binding (**Fig. S2F–J**), indicating that they disrupt distinct steps of the interaction. Together, these mutations impair both the initial docking and the final stabilisation of the VBS–VD1 complex. Consistent with these results, independent fluorescence imaging experiments showed strongly reduced FA enrichment of VD1 and full-length vinculin carrying the I12K/A50I mutations (**Fig. S8**).

### Reconsidering talin-independent effects inferred from the A50I mutant

Because A50I vinculin retains measurable binding to talin, experiments using this construct represent partial disruption of the talin–vinculin interaction rather than its complete loss. The I12K/A50I mutant introduced here provides a tool to uncouple vinculin function from talin binding and directly test talin-dependent vinculin regulation in cells. Using this mutant, we show that vinculin localisation to FAs is highly influenced by talin binding, but not solely dependent upon it. Moreover, we find that talin-vinculin interactions are not required for release of vinculin’s head-tail inhibition and activation. Perhaps most interestingly, we find that vinculin can still experience mechanical tension in focal adhesions even when talin-binding is abolished. These findings indicate that talin functions not simply as a mechanical linker but as a spatial organiser of force transmission within adhesion complexes, and that vinculin can be recruited, activated, and mechanically loaded through multiple, parallel molecular pathways.

In addition to broad functional analysis, this construct was also used to assess the role of talin in the spatial regulation of vinculin function. For instance, these experiments revealed that cell-level spatial regulation of vinculin activation requires talin interactions, while such tuning of vinculin load does not. This is consistent with the observation that regulation of vinculin activation and loading are separable, and potentially important in the regulation of focal adhesion reinforcement during migration (Grashoff et al. 2010; Hernández-Varas et al. 2015). Such insights went previously unnoticed with the A50I vinculin construct, potentially due to the residual talin-binding sites masking other signals in the cell periphery. Moreover, analysis of vinculin load at individual focal adhesions using the talin-binding-null mutant revealed that the vinculin-talin interaction is required for FA-level vinculin tension gradients (**Fig. 6F-G**). Specifically, talin binding enables peak vinculin tension near membrane-proximal adhesion edges. While loss of FA-level tension gradients was previously observed with A50I vinculin TS (Rothenberg et al. 2018), observation of the loss of loading gradient with the I12K/A50I vinculin TS unambiguously establishes the role of the vinculin-talin interaction. Interestingly, other work has also revealed that vinculin tension and traction force are highly correlated in the peripheral FAs of cells (Aytekin et al. 2026). Together with our results, this suggests that a primary role of the talin-vinculin interaction may be facilitating the generation of large traction forces at mature FAs.

Overall, these results indicate that re-examining these processes using a talin-binding-null vinculin will clarify how mechanical signals are transmitted and integrated within adhesion complexes. More broadly, they show that force transmission through adhesion complexes is distributed across multiple interacting pathways rather than being mediated solely through the talin–vinculin interaction.

## Materials and Methods

### Genetic screen for mutations that suppress VD1 lethality in *Drosophila*

Expression of chicken vinculin VD1 in flies using the conditional Gal4 system results in lethality but does not cause cytoplasmic aggregation of the integrin machinery observed with expression of *Drosophila* VD1 (Maartens et al. 2016). Chicken VD1 was therefore used for the suppressor screen because it caused pupal lethality without the cytoplasmic aggregation phenotype observed with *Drosophila* VD1 (Maartens et al. 2016). For the screen, we used the C57 Gal4 driver (FBti0016293), which is expressed primarily in the muscle, and when crossed with UAS::ChickenVD1-RFP (FBtp0116463) resulted in pupal lethality (<0.1% escapers). Males homozygous for UAS::ChickenVD1-RFP were treated with 25 mM EMS and crossed to w; CyO/Gla; P{w+,Gal4}C57 virgin females in 30 bottles each containing 25 females and 12 males (day 1). Flies were transferred to new bottles on day 2,3,5, and 7, with males removed on day 3. Dead pupae were counted in a couple of bottles from each set to estimate total number screened.

From approximately 66,000 flies screened, 70 adult flies were recovered, 47 of which were fertile, and 20 retested. Of these, 18 had mutations in the VD1 construct with 3 having large deletions and 15 having single base changes: 7 premature stops, 4 missense mutations in the RFP sequence and 4 missense mutations in VD1.

The expression of the VD1 missense mutants, when driven with C57-Gal4 in muscles and examined in first instar larvae by confocal microscopy, appeared normal; all were still recruited to the sites of integrin adhesion where the muscle attaches to the tendon matrix, at similar levels to the wild-type construct.

### Plasmids and Peptides

Plasmids encoding VD1 (2–258) in pET151D-TOPO were synthesised by GeneArt based on the mouse vinculin sequence (UniProt: P18206). Constructs included WT, A50I, I12K, P15L, I12K/A50I and P15L/A50I.

Mouse talin-1 and α-catenin VBS peptides were synthesised by GLBiochem (Shanghai) with a non-native cysteine added at either the N- or C-terminus (for fluorescent labelling in FP assays) as indicated.

Talin H33 (C1516-1549): C-TANPTAKRQFVQSAKEVANSTANLVKTIKALDGD

Talin H46 (C1943-1969): C-DVYTKKELIESARRVSEKVSHVLAALQ

Talin H50 (C2072-2103): C-EDPETQVVLINAVKDVAKALGDLISATKAAAG

Talin H58 (2345-2369C): ILEAAKSIAAATSALVKAASAAQRE-C

α-catenin VBS (C326-355): C-RDDRRERIVAECNAVRQALQDLLSEYMGNA

### Protein expression and purification

BL21(DE3) Star cells transformed with VD1 constructs were grown in LB supplemented with 100 µg/mL ampicillin. Protein expression was induced with 400 μM IPTG at OD_600_ ∼0.6-0.8 and cultures were incubated overnight at 20°C. Cells were harvested and resuspended in lysis buffer (50 mM Tris pH 8, 250 mM NaCl, 10% w/v glycerol) at 5 mL per gram of wet pellet. Cells were lysed by sonication (3 s pulses at 35% amplitude with 7 s off-intervals; for 2 minutes total sonication time). Cellular debris was removed by centrifugation (40,000 x g, 45 min, 4°C). The supernatant was filtered (0.22 μm) and loaded onto a 5 mL HisTrap column (Cytiva) using an ÄKTA Start system. Bound proteins were eluted with a 0–300 mM imidazole gradient. Fractions containing protein were identified by SDS–PAGE, pooled, and diluted into 20 mM Tris pH 8.0 prior to loading onto a 5 mL HiTrap Q HP anion-exchange column (Cytiva). Proteins were eluted using a 0–750 mM NaCl gradient. Selected fractions were combined and dialysed overnight at 4°C against PBS (137 mM NaCl, 2.7 mM KCl, 10 mM Na_2_HPO_4_, 1.8 mM KH_2_PO_4_, pH 7.4) using 10 kDa MWCO SnakeSkin dialysis tubing (Thermo Fisher Scientific). Protein concentrations were determined using a NanoDrop 2000c (Thermo Fisher Scientific), and samples were concentrated to 350-450 μM using 10 kDa MWCO centrifugation filters. Aliquoted protein samples were flash-frozen in liquid nitrogen and stored at −20°C.

### Biophysical characterisation of VD1 mutants

#### Fluorescence Polarisation (FP)

Binding affinities of talin and α-catenin VBS peptides to WT and mutant VD1 were measured using fluorescence polarisation. VBS peptides were prepared as 2.5 mM stocks in PBS. The synthetic peptides were coupled to a thiol-reactive fluorescein fluorophore (Thermo Fisher Scientific) via the terminal non-native cysteine (Vicente-Manzanares 2021). Uncoupled dye was removed using a PD-10 desalting column.

Purified VD1 stocks were prepared at 80 μM (2x final concentration) in PBS supplemented with 1 mM TCEP, and fluorescein-labelled VBS peptides were prepared at 1 μM (2x final concentration) in PBS containing 0.02% v/v Tween-20. Serial 1:2 dilutions of VD1 mutants were prepared in triplicate in black 96-well plates (Nunc) with a final volume of 50 μL per well. Fluorescein-labelled peptide (50 µL) was added to each well, yielding a final peptide concentration of 0.5 μM and a serial dilution series of VD1 starting at 40 μM. Fluorescence polarisation was measured at room temperature using a Hidex plate reader. Data were analysed using GraphPad Prism. Dissociation constants (K_d_) were obtained by fitting the data to a one-site total binding model.

#### Circular Dichroism (CD)

Circular dichroism spectroscopy was used to assess the secondary structure and thermal stability of WT and mutant VD1 proteins. Samples were prepared at a final concentration of 12 μM either alone or with 12 μM VBS peptide in PBS supplemented with 1 mM TCEP and analysed in a QS 0100 quartz cuvette. CD spectra were recorded using a Jasco J-1100 spectrometer between 198- 260 nm at 20 and 90°C. Thermal denaturation experiments were performed by monitoring ellipticity at 222 nm while increasing the temperature from 20-90°C in 0.5°C increments. Melt curves were analysed in GraphPad Prism and fitted with a biphasic model to determine the melting temperatures (T_m_) of the two VD1 subdomains.

#### GST pulldown assays

GST-talin rod was expressed in *E. coli* BL21(DE3) Star cells as described above. Following sonication, the lysate was loaded onto a 5 mL GSTrap column (Cytiva) using an ÄKTA Start system. The column was washed with wash buffer (50 mM Tris pH 8, 600 mM NaCl, 30 mM imidazole, 10% w/v glycerol, 0.2% v/v Triton X-100, 1 mM TCEP) followed by PBS. GST-talin rod was eluted with PBS containing 10 mM L-glutathione, diluted 1:10 in PBS to reduce the glutathione concentration and concentrated to 50 μM using a 30 kDa MWCO micro-concentrator.

GST-talin and VD1 (WT and mutants) were diluted to 10 and 50 μM, respectively, in PBS supplemented with 1 mM TCEP. Equal volumes were mixed to yield final concentrations of 5 μM GST–talin and 25 μM VD1 (1:5 molar ratio) and incubated for 1 h at room temperature with agitation. For pulldown assays 50 μL of GST-talin/VD1 was added to 50 μL of a 50% v/v slurry of glutathione agarose beads (Thermo Fisher Scientific) in PBS containing 1 mM TCEP and incubated for 1 h at room temperature with agitation. Control samples were prepared by diluting 50 μL of the GST-talin/VD1 mixture with 50 μL PBS containing 1 mM TCEP. The beads were pelleted by centrifugation (6,000 x g, 1 min), washed twice with PBS containing 1 mM TCEP, and analysed by SDS-PAGE using a 10% polyacrylamide gel. Gels were stained with Coomassie Brilliant Blue and imaged with a Bio-Rad ChemiDoc MP imaging system (590/110 white transmission).

#### Crystallisation and structure determination

The structure of A50I VD1 in complex with talin H50 was determined by X-ray crystallography. Purified A50I VD1 was incubated overnight at 4°C in 20 mM Tris pH 8, 150 mM NaCl with TEV protease (40:1) to remove the N-terminal His-tag. TEV protease and the cleaved His-tag were removed using a HisTrap nickel affinity column (Cytiva). The cleaved protein was concentrated to 18 mg/mL, flash frozen in liquid nitrogen, and stored at −20°C.

A50I VD1 was mixed with H50 peptide at a 1:1.1 molar ratio to a final concentration of 7.5 mg/mL in 20 mM Tris pH 8, 150 mM NaCl, 1 mM DTT. Hanging-drop vapour diffusion crystallisation trials were set up by mixing protein solution and mother liquor (100 mM bis-tris pH 6.0, 15% PEG 3350) in a 1:1 ratio (2 μL total drop volume) and equilibrating over a 600 μL reservoir. Crystals were harvested and flash-frozen in liquid nitrogen in mother liquor supplemented with 30% glycerol as cryo-protectant.

X-ray diffraction was collected remotely at beamline I24 (Diamond Light Source, UK) using a DECTRIS EIGER2 9M detector at a wavelength of 0.8 Å. Diffraction data were indexed, integrated and scaled using xia2 (3dii pipeline). The structure was solved by molecular replacement with MOLREP (Vagin and Teplyakov 2010) using the structure of WT VD1-H50 structure (PDB ID: 4DJ9 (Yogesha et al. 2012)) as the search model. Iterative rounds of real and reciprocal space refinement were performed using Coot (Emsley et al. 2010) and REFMAC (Murshudov et al. 2011) within the CCP4i2 suite (Winn et al. 2011), with final refinement carried out in Phenix (Liebschner et al. 2019). Data collection and refinement statistics are reported in Table 1. Structural figures were prepared using PyMOL (Schrodinger, LLC).

**Table 1:** X-ray data collection and refinement statistics for A50I VD1 (2-258) in complex with talin-1 H50.

|  |  |
| --- | --- |
| Synchrotron and Beamline | Diamond I24 |
| Space group | P2<sub>1</sub>2<sub>1</sub>2<sub>1</sub> |
| Molecule/ASU | 1 |
| Cell dimensions |  |
| *a*, *b*, *c* (Å) | 43.7, 75.8, 82.58 |
| $\alpha$, $\beta$, $\gamma$ (°) | 90, 90, 90 |
| Resolution (Å) | 38.63 – 1.49 (1.52 – 1.49) |
| *R*<sub>merge</sub> | 0.078 (1.95) |
| *I* / $\sigma$ | 10.1 (0.7) |
| CC(1/2) | 0.999 (0.557) |
| Completeness (%) | 99.7 (99.6) |
| Redundancy | 7.4 (6.7) |
| Refinement |  |
| Resolution (Å) | 38.63-1.49 |
| No. reflections | 45809 (2689) |
| *R*<sub>work</sub> / *R*<sub>free</sub> | 0.1895/0.2350 |
| No. atoms |  |
| Protein | 2610 |
| Water | 230 |
| *B*-factors (Å<sup>2</sup>) |  |
| Protein | 24.4 |
| Water | 52.8 |
| R.M.S. deviations |  |
| Bond lengths (Å) | 0.005 |
| Bond angles (°) | 0.680 |
| Ramachandran plot |  |
| Favoured/outlier (%) | 98.89/0.37 |
| Rotamer |  |
| Favoured/poor (%) | 94.72/0.75 |
| MolProbity scores |  |
| Clash score all atoms | 5.00 |
| PDB accession no. | 9SR0 |

Data collected from a single crystal.
Values in parentheses are for highest-resolution shell.

### Single-Molecule Stretching

Single-molecule manipulation experiments were performed as previously described (Yao et al. 2016), using a custom-built high-force magnetic tweezers instrument capable of applying forces up to 100 pN with an extension resolution of ∼1 nm (Hu Chen et al. 2011; Zhao et al. 2017). Full-length talin rod constructs (R1–R13) were immobilized on glass coverslips within a laminar flow chamber via HaloTag–ligand chemistry and tethered to 2.8 µm paramagnetic beads through biotin–streptavidin interactions. To preserve the native folded state of the R1–R13 construct during solution exchange of the vinculin head domain (VD1), a buffer-isolation membrane well array was employed (Le et al. 2015). All measurements were conducted at a constant loading rate of 2 pN/s. Force calibration was performed using a standard calibration curve (Zhao et al. 2017), requiring a single reference point correlating force with magnet–bead distance. The well-characterized DNA overstretching transition (Smith et al. 1996; Cohen et al. 2005) was used as the reference, yielding an overall force calibration uncertainty of less than 5% (Zhao et al. 2017).

### Tension sensor and FRET studies

FRET conformation and tension sensors were used to quantify vinculin conformation and mechanical loading across vinculin in focal adhesions.

#### Vinculin Tension Sensor I12K Site-Directed Mutagenesis

VinTS was a gift from Martin Schwartz (Addgene plasmid # 26019; http://n2t.net/addgene:26019; RRID:Addgene_26019) (Grashoff et al. 2010). VinTS-A50I (Addgene plasmid # 111827; http://n2t.net/addgene:111827; RRID:Addgene_111827), VinTS-I997A (Addgene plasmid # 111828; http://n2t.net/addgene:111828; RRID:Addgene_111828), VinTS-A50I/I997A (Addgene plasmid # 111829; http://n2t.net/addgene:111829; RRID:Addgene_111829), and VinCS (Addgene plasmid # 200309; http://n2t.net/addgene:200309; RRID:Addgene_200309) were described previously (Rothenberg et al. 2018).

The I12K mutation was introduced into VinTS-A50I and VinTS-A50I/I977A plasmids using site-directed mutagenesis with overlapping primers Forward: GCACCATCGAGAGCAAATTGGAGCCCGTGGC, Reverse:

GCCACGGGCTCCAATTTGCTCTCGATGGTGC. Whole plasmid sequencing, performed by Plasmidsaurus (London, UK) using Oxford Nanopore technology with custom analysis and annotation, was used to sequence verify isolated plasmids.

#### Generation of vinculin mutant constructs

Construction of pcDNA3.1-VinTS, pcDNA3.1-VinCS, and pcDNA3.1-VinTS A50I, and corresponding cell lines where applicable, have been described previously (Grashoff et al. 2010; Rothenberg et al. 2018).

To create a vinculin conformation sensor harbouring the A50I mutation, the vinculin head fragment containing this mutation was isolated from pcDNA3.1-VinTS A50I using *5’-HindIII / 3’-BamHI*. This DNA fragment was then inserted into pcDNA3.1-VinCS digested with *5’-HindIII / 3’-BamHI*. Similarly, the vinculin I12K/A50I conformation sensor was created by digesting pcDNA3.1- VinTS I12K/A50I with *5’-NheI / 3’-BamHI* and inserting this fragment into pcDNA3.1-VinCS digested with *5’-NheI / 3’-BamHI*. An intermediate pcDNA3.1-VinCS I997A construct was created by PCR-based mutagenesis on the pcDNA3.1-VinCS backbone, using the following primer pairs: 5’-GCA GCT CAA AGC TCT TTC CAC AGT GAA AGC TAC CAT GCT G-3’ and 5’-AGG AGC TAA CCG CTT TTT TGC ACA A-3’; 5’-TTG TGC AAA AAA GCG GTT AGC TCC T-3’ and 5’-CTG TGG AAA GAG CTT TGA GCT GCG TGC TGA TGG TTG-3’. The PCR fragments were joined through Gibson assembly. This intermediate construct was digested using *5’-NheI / 3’-BamHI*. The *5’-NheI / 3’-BamHI* digested fragment from pcDNA3.1-VinTS I12K/A50I was then inserted to create the vinculin I12K/A50I/I997A conformation sensor.

#### Cell culture and transfection

Vinculin (-/-) Mouse Embryonic Fibroblasts (MEFs) and vinculin (-/-) MEFs stably expressing VinTS or VinCS constructs were maintained in high glucose Dulbecco’s Modified Eagle’s Medium containing L-glutamine, sodium pyruvate, and sodium bicarbonate (Sigma Aldrich) supplemented with 10% (v/v) Fetal bovine serum (Cytiva), 1% (v/v) penicillin-streptomycin (Thermo Fisher), and 1% (v/v) Minimum Essential Medium Non-Essential Amino Acids (Gibco). Cells were passaged every 2 days using 0.05% Trypsin-EDTA (Gibco). All cell lines were maintained at 37 °C with 5% CO2 perfusion.

To express the vinculin mutant TS and CS constructs, vinculin (-/-) MEFs were transfected at 60-70% confluency in 6-well tissue culture plates using Lipofectamine 2000 (Thermo Fisher), according to manufacturer protocol.

#### Cell seeding and fixation

For imaging, 6-well glass bottom dishes, with no. 1.5 cover glass (CellVis) were functionalised with 10 µg/ml fibronectin (Sigma-Aldrich) diluted in PBS overnight at 4°C. Wells were rinsed with PBS prior to cell seeding. Approximately 100,000 cells were plated per well. Cells were allowed to spread in cell culture media for 4-6 hours prior to fixation.

Immediately prior to fixation, cells were rinsed twice with PBS. Cells were then incubated for 10 minutes in 4% (v/v) methanol-free paraformaldehyde (Electron Microscopy Sciences) in PBS. Post fixation, cells were rinsed twice with PBS and left immersed in PBS for imaging.

#### FRET Imaging

An Olympus IX83 inverted epifluorescent microscope (Olympus), equipped with a LambdaLS 300 W ozone-free xenon bulb (Sutter Instruments) and sCMOS ORCA-Flash4.0 V2 camera (Hamamatsu Photonics), was used to image samples. Images were acquired at 60x magnification (UPlanSApo 60X/NA1.35 Objective, Olympus). MetaMorph Advanced software (Olympus) was used to control image acquisition.

FRET images were acquired with a three-image sensitised emission acquisition sequence (Huanmian Chen et al. 2006). To image the mTFP1-Venus sensors a custom filter set was used, comprised of an mTFP1 excitation filter (ET450/30x; Chroma Technology Corp), mTFP1 emission filter (FF02-485/20-25; Semrock), Venus excitation filter (ET524/10x; Chroma Technology Corp), Venus emission filter (FF01-571/72; Semrock), and dichroic mirror (T450/514rpc; Chroma Technology Corp). Images were acquired sequentially across the acceptor channel (Venus excitation, Venus emission, 500 ms exposure), FRET channel (mTFP1 excitation, Venus emission, 750 ms exposure), and donor channel (mTFP1 excitation, mTFP1 emission, 750 ms exposure).

#### Quantitative FRET efficiency calculations from sensitised emission

FRET efficiency was calculated on a pixel-by-pixel basis using custom MATLAB code as described previously (Rothenberg et al. 2018; Gates et al. 2019). Prior to analysis, images were corrected for dark current, uneven illumination, background intensity, and registered.

Spectral bleed-through coefficients were determined using donor-only and acceptor-only samples. Donor bleed-through (dbt) was calculated as:

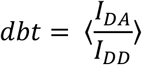

where *I_DA_* is the FRET channel intensity and *I_DD_* is the donor channel intensity. Acceptor bleed-through coefficients (abt) were, similarly, calculated for Venus as:

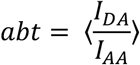

where *I_AA_* is the intensity in the acceptor channel and data were binned by acceptor channel intensity. Corrected FRET intensity was calculated as:

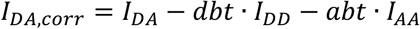

where *I_DA,corr_* is the corrected FRET intensity, also denoted by *F_C_* in the literature.

To calculate FRET efficiency, the proportionality constant, *G*, was determined after imaging donor-acceptor fusion constructs of differing, but constant, FRET efficiencies. *G* is a constant describing the increase in acceptor intensity, due to sensitised emission, relative to the decrease in donor intensity, due to quenching (Huanmian Chen et al. 2006; Gates et al. 2019) and is calculated as:

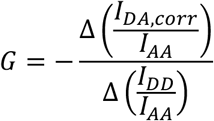

where *Δ* indicates the change between two donor-acceptor fusion constructs. FRET efficiency (*E*) can then be calculated as:

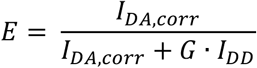

To calculate FRET-PAIR stoichiometry, a separate proportionality constant, *k*, is determined from donor-acceptor fusion construct imaging. *k* is a constant that describes the ratio of donor to acceptor fluorescence intensity for equimolar concentrations of the fluorophores in the absence of FRET (Huanmian Chen et al. 2006; Gates et al. 2019) and is calculated as:

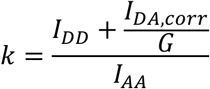

FP stoichiometry (S) (Coullomb et al. 2020) can then be calculated as:

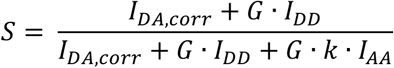

#### Image Segmentation, FA Analysis, and Data Selection Criteria

Segmentation was performed on acceptor-channel images, which are independent of FRET and proportional to vinculin concentration. Segmentation of focal adhesions was performed using a previously described water-based algorithm (Rothenberg et al. 2018). The output of FA segmentation was a binary mask, which was inverted to measure cytosolic parameters. Single cells were identified based on closed boundaries, drawn by the user, on unmasked acceptor channel images. These single cell masks were used to calculate cell regions based on the Euclidean distance from the cell centre, with the inner region defined as distances within the 0-20^th^ percentile from cell centre, middle region as 20-50^th^ percentile, and outer region as 50-100^th^ percentile. Cells that were not fully spread, indicated by a cell area less than 23.3 µm^2^ or with less than 10 focal adhesions, were discarded.

Constructs that were expressed via transient transfection were expression matched to the stably expressed vinculin constructs, which have been previously sorted to match physiological vinculin levels. Specifically, acceptor intensity was used as a proxy for expression and cells with an average acceptor intensity below a threshold value of 1000 were excluded. To ensure proper functioning of the FRET sensor, cells with an average FP stoichiometry below 0.33 and above 0.67 were excluded from downstream analysis.

Line scans of single focal adhesions were performed in ImageJ software (National Institutes of Health). Specifically, the line tool was used to draw a line axially, starting at the tip distal to the cell body, along single, large FAs in the cell periphery. This line was used to visualise the acceptor intensity and corresponding FRET efficiency profiles. The profiles were graphed in MATLAB, and a cubic spline-based smoothing algorithm was used for noise reduction.

### Statistical Analysis

Statistical analyses were performed using JMP Pro software (SAS). All experimental data sets were initially assessed for unequal variances using Levene’s. The multi-sample datasets were then assessed using a one-way Welch’s ANOVA followed by post hoc Steel-Dwass test, when necessary (unequal variances). A value of p < 0.05 was considered statistically significant. In figures, groups not connected by the same letter are statistically different. Box plots were created using GraphPad software (Dotmatics). Box depicts the median as the centre value, the 25^th^ percentile as the lower bound, and the 75^th^ percentile as the upper bound. The whiskers extend to the minimum and maximum of the data.

## Data Accessibility

The atomic coordinates and structure factors for the A50I VD1–H50 crystal structure have been deposited in the Protein Data Bank under accession code **9SR0**. Plasmids encoding I12K/A50I vinculin constructs have been deposited at Addgene and are available to the community (www.addgene.org/ben_goult).

## Acknowledgements

This work was supported by a Cancer Research UK Programme Grant (CRUK-A21671), a British Heart Foundation Special Project Grant (SP/F/23/150045), the Research Council of Finland (363941), and Sigrid Jusélius Foundation and Cancer Foundation Finland. We thank Diamond Light Source, UK, for synchrotron access (MX15591 BAG) and Dr Svetlana Antonyuk (Institute of Systems, Molecular and Integrative Biology, University of Liverpool) for assistance with crystal production and storage, X-ray diffraction data collection, and generation of the initial structural model. We thank Daniel Cooper for critical reading of the manuscript. We acknowledge Biocenter Finland and Tampere imaging facility for infrastructure support and the CSC (Finnish IT Center of Science).

## Supplemental Information for

Supplementary figures provide structural rationale for the mutant design, validation of VD1 folding and binding properties, and additional controls for the single-molecule experiments

**Figure S1.**
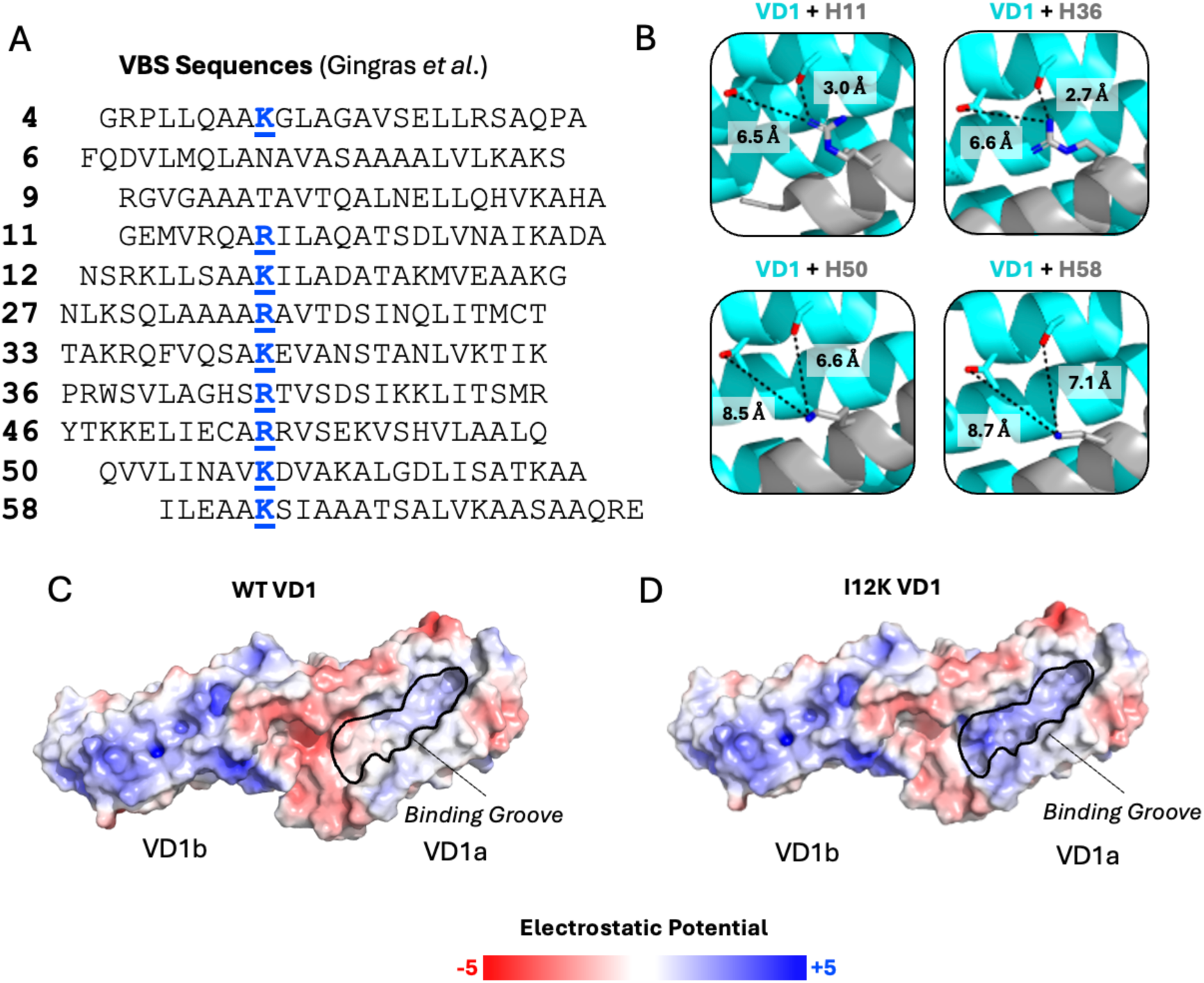
Structural rationale for the I12K/A50I mutation related to Fig. 3. **(A)** Talin-1 VBS sequence alignment from (Gingras et al. 2005), showing the position of a highly conserved positively charged residue (Arg, Lys) highlighted in blue. **(B)** Crystal structures of VD1 bound to H11 (PDB ID: 1ZVZ) (Gingras et al. 2005), H36 (PDB ID: 1ZW3) (Gingras et al. 2005), H50 (PDB ID: 4DJ9) (Yogesha et al. 2012), and H58 (PDB ID: 1ZW2) (Gingras et al. 2005) showing that the conserved positively charged residue on each VBS is positioned adjacent to serine or threonine residues in VD1. Estimated distances between VD1 Ser/Thr and VBS Arg/Lys are indicated (Å). **(C)** APBS electrostatic surface representations of WT and I12K VD1 in the bound state showing that the I12K substitution increases positive electrostatic potential within the VBS-binding groove.

**Figure S2.**
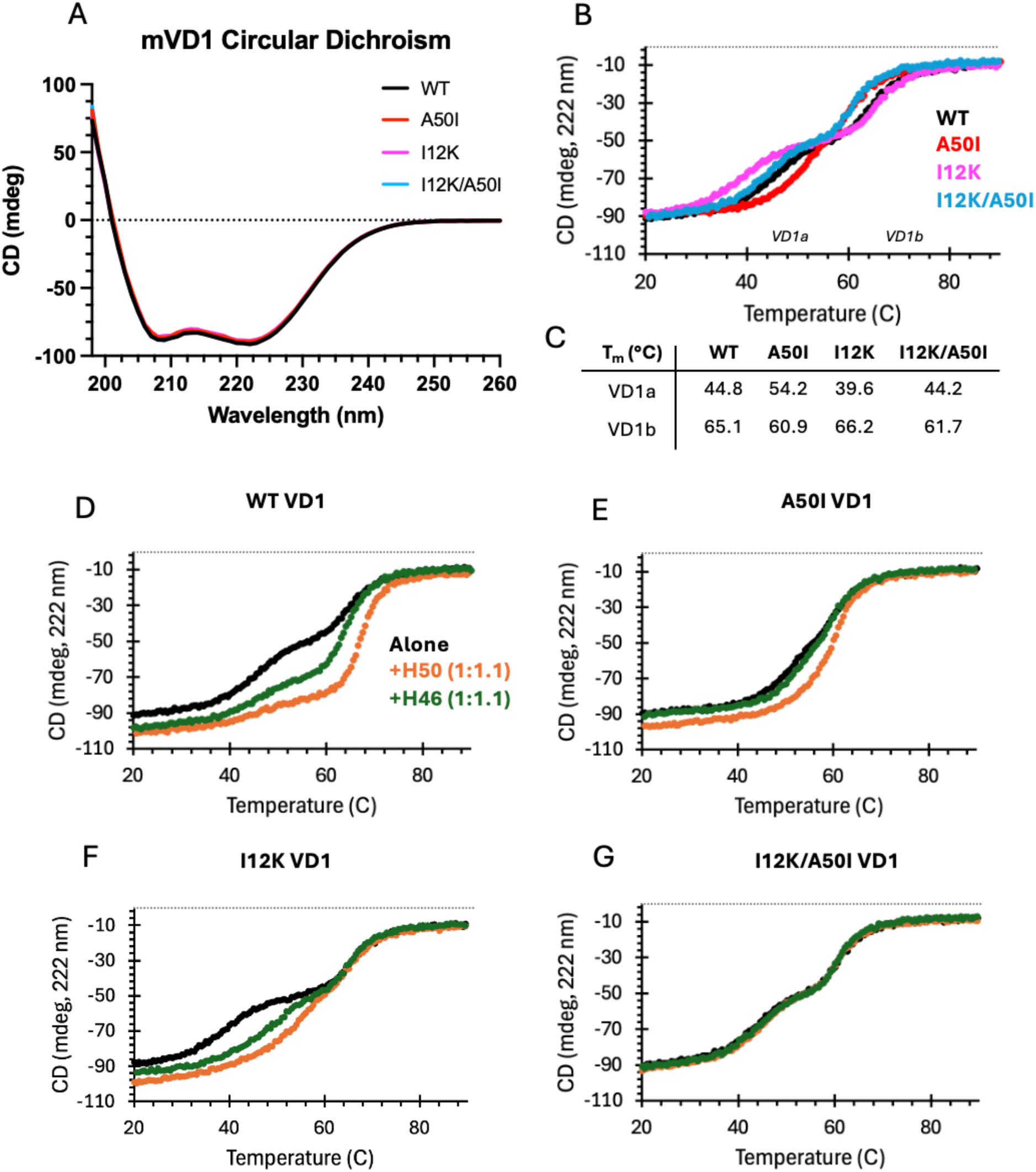
Circular dichroism analysis of VD1 mutants stability related to Fig. 4. **(A)** CD spectra of WT, A50I, I12K, and I12K/A50I VD1 (12 µM). **(B)** CD thermal stability profiles (20-90 °C) of VD1 WT and mutants showing melting of VD1a and VD1b subdomains. (**C**) Melting temperatures (T_m_) corresponding to VD1a and VD1b transitions determined by biphasic fitting in GraphPad Prism. **(D-G)** Thermal denaturation profiles monitored by CD at 222 nm of (**D**) WT, (**E**) A50I, (**F**) I12K, or (**G**) I12K/A50I VD1 alone (black) or in complex (1:1.1) with H50 (orange) or H46 (green).

**Figure S3.**
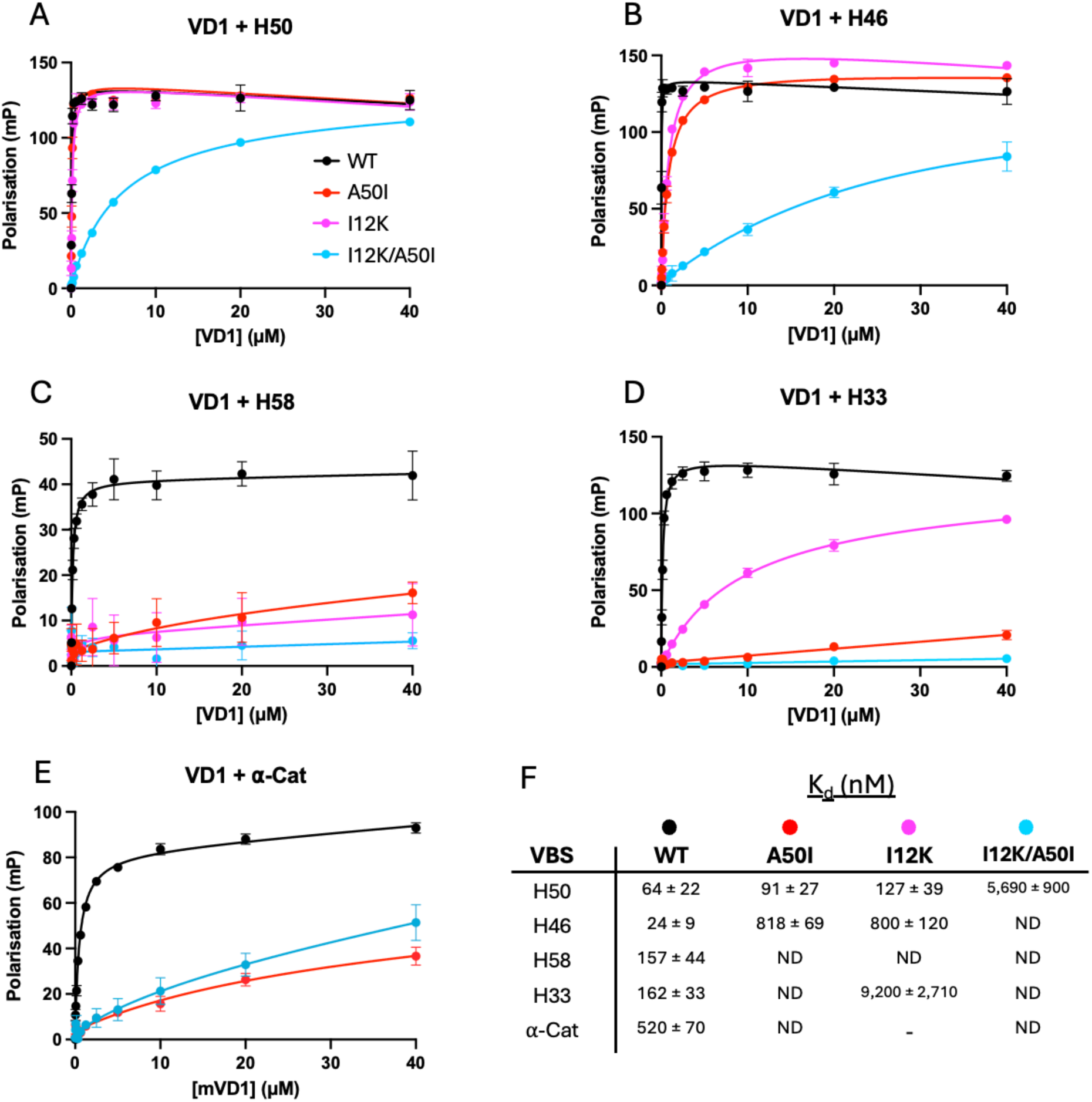
Fluorescence polarisation analysis of VD1 mutant binding related to Fig. 1 and Fig. 4. Fluorescence polarisation binding curves comparing WT (black), I12K (magenta), A50I (red), and I12K/A50I (blue) VD1 interactions with **(A)** H50, **(B)** H46, **(C)** H58, **(D)** H33, and **(E)** the α-catenin VBS.

**Figure S4.**
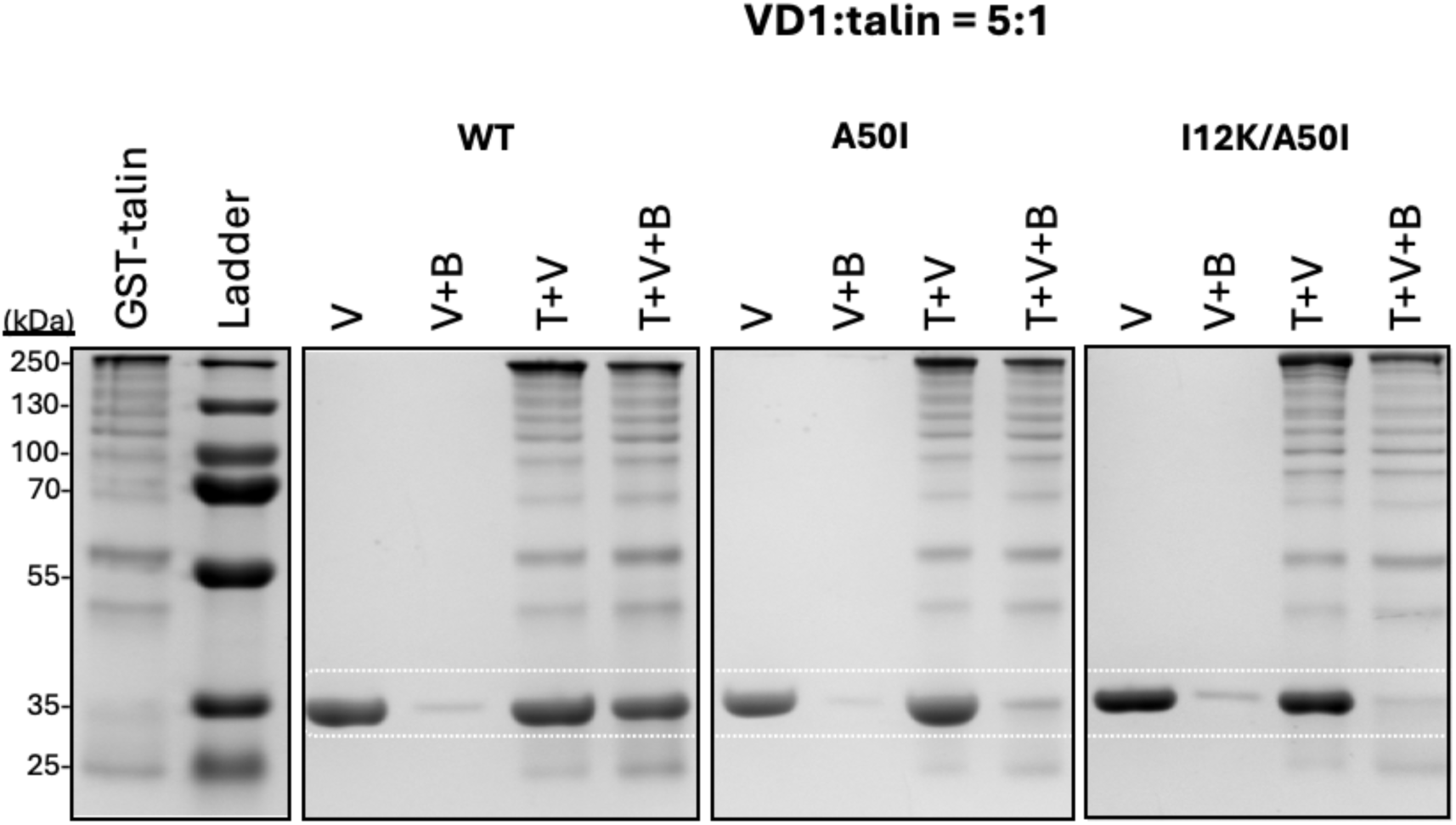
GST pulldown analysis of VD1 binding to the talin rod. Representative SDS-PAGE gels of GST pulldown assays using GST–talin-1 rod and WT, A50I, or I12K/A50I VD1 (5:1 VD1:talin molar ratio). Lanes (left to right): GST–talin + beads, protein ladder, VD1 alone, VD1 + beads control, and VD1 + GST–talin in the absence or presence of beads. VD1 bands are highlighted by dashed white boxes.

**Figure S5.**
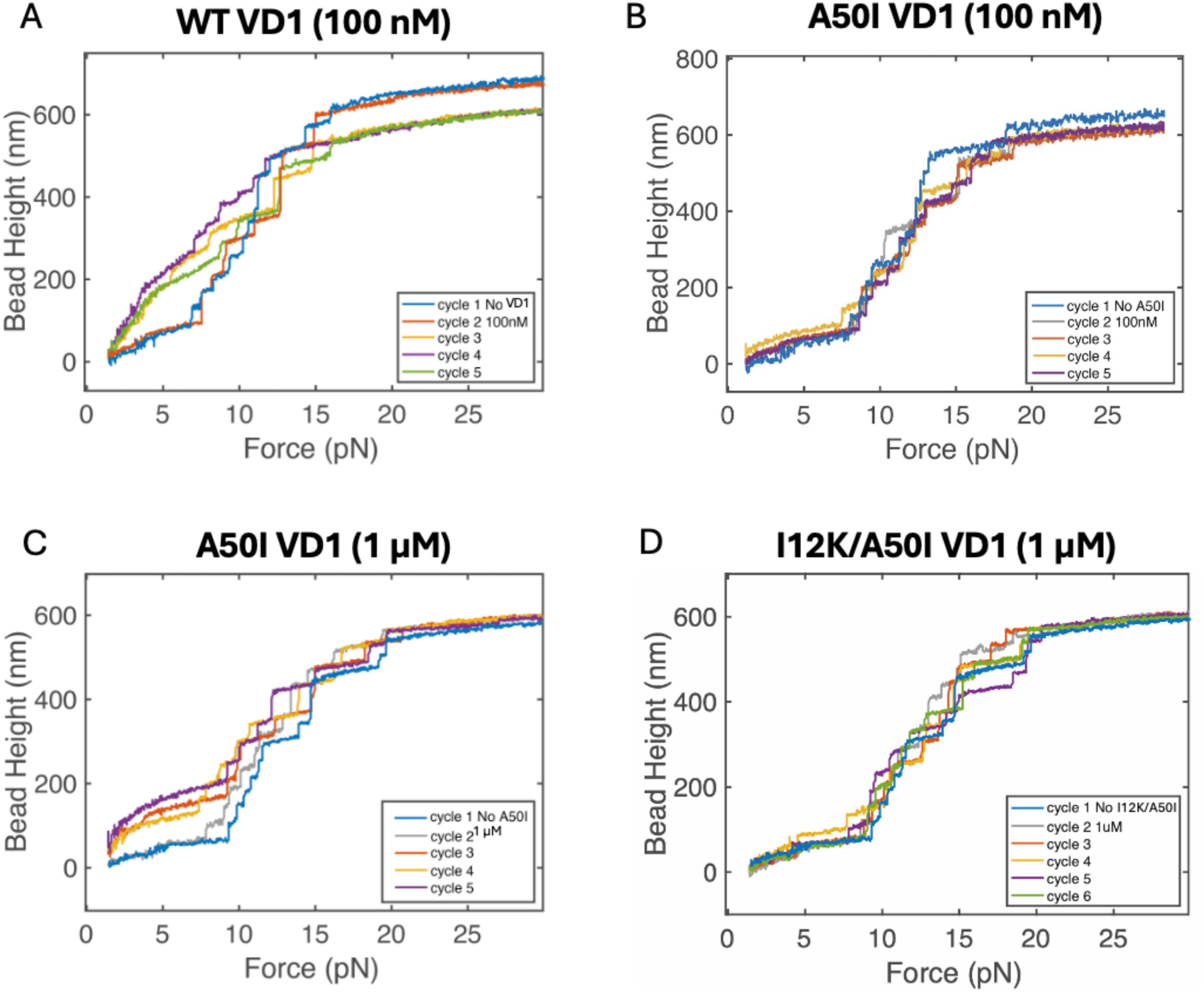
Single-molecule stretching experiments related to Fig. 5. **(A-D)** Representative unfolding force–extension curves of the talin-1 rod recorded at a loading rate of 2 pN s⁻¹ in the presence of **(A)** WT VD1 (100 nM), **(B)** A50I VD1 (100 nM), **(C)** A50I VD1 (1 µM), or **(D)** I12K/A50I VD1 (1 µM).

**Figure S6.**
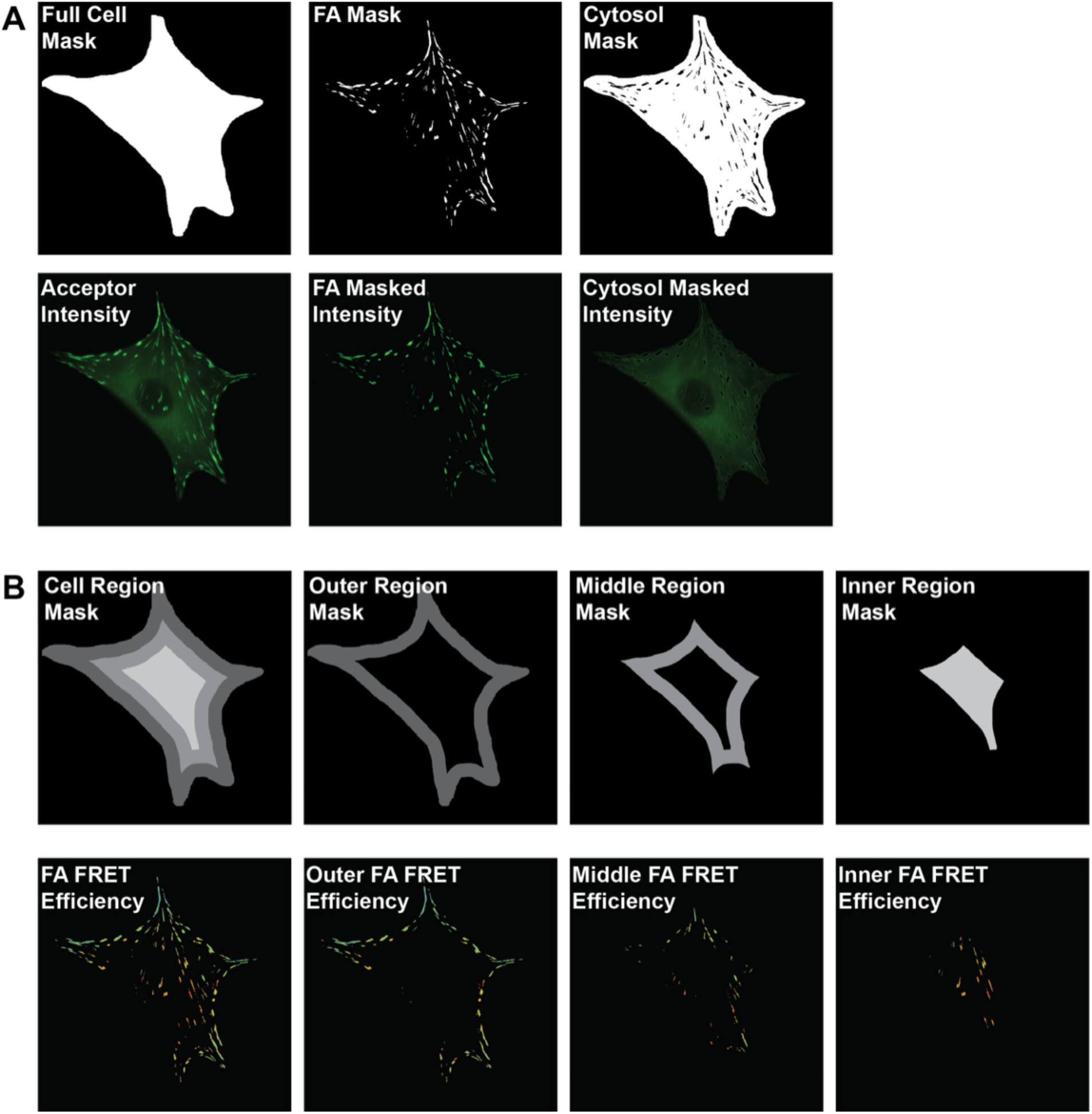
Localisation and region-specific FRET analysis for vinculin tension and conformation sensors. **(A)** Representative masks (top row) and corresponding acceptor intensity images (bottom row) used for cytosol-to-FA intensity ratio analysis. Focal adhesion segmentation was performed using a previously described custom MATLAB algorithm. **(B)** Representative cell region masks (top row) and corresponding FA-masked FRET efficiency images (bottom row) used for region-specific FRET analysis. Regions were defined as the 100^th^ to 50^th^, 50^th^ to 20^th^, and 20^th^ to 0 percentiles of Euclidean distance from the cell centre, corresponding approximately to outer, middle, and inner cell regions.

**Figure S7.**
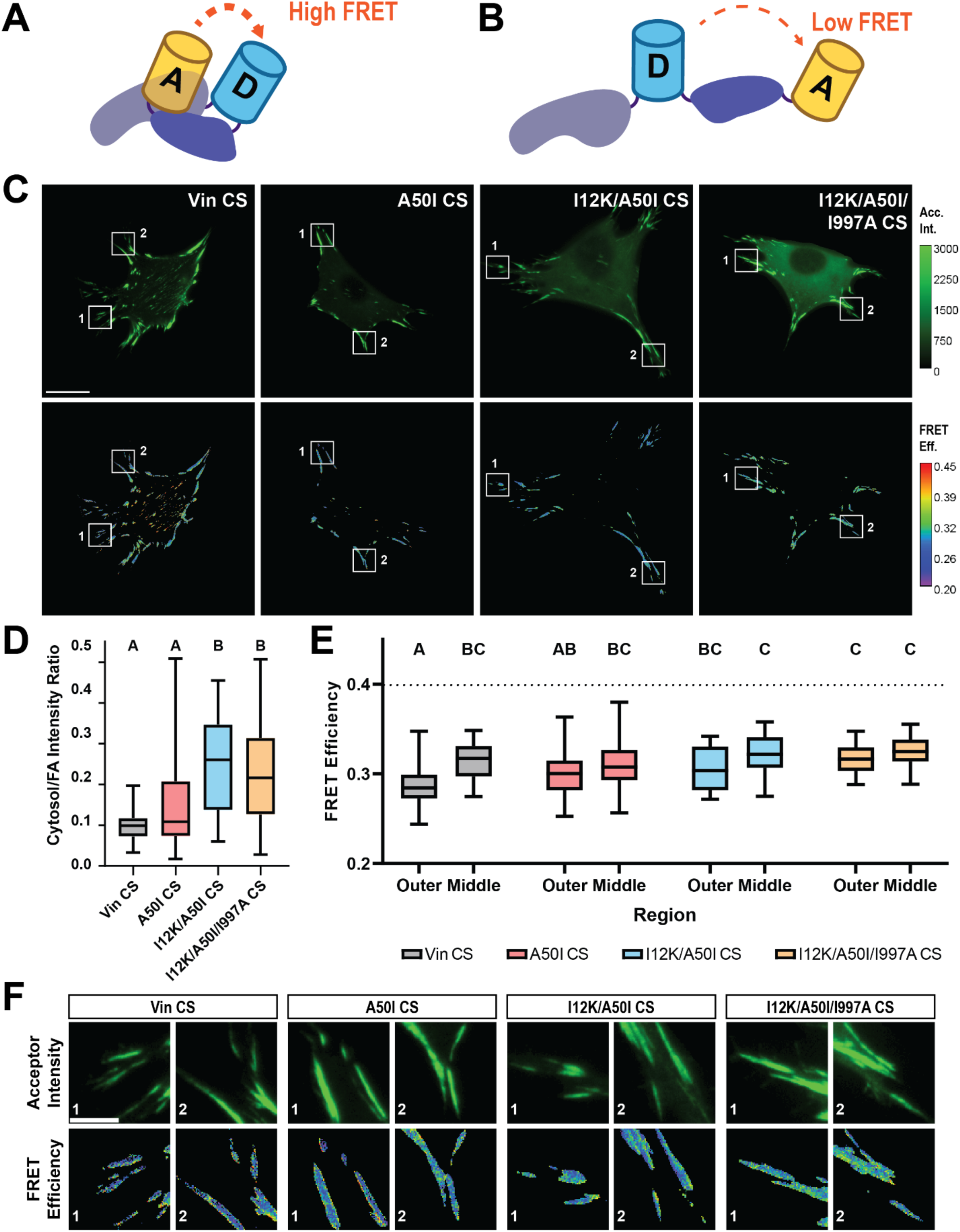
Vinculin conformation is minimally affected by talin and actin binding at focal adhesions. **(A-B)** Schematic diagrams of the vinculin conformation sensor in **(A)** inactive (high FRET) and **(B)** active (low FRET) conformations. **(C)** Representative images of acceptor (top row) and masked FRET efficiency (bottom row) for vinculin (-/-) MEFs expressing vinculin conformation sensor constructs. Scale bar, 20 µm. Box and whisker plots are shown for **(D)** cell-averaged cytosol to focal adhesion acceptor intensity ratios and **(E)** cell-averaged FRET efficiency within outer and middle regions of the cell for each vinculin CS construct. Letters above box plots represent statistical significance, with box plots with no letter in common being significantly different from one another at p < 0.05. One-way ANOVA with Steel–Dwass multiple comparisons test was used for the statistical analysis. Dashed line represents the previously established closed FRET efficiency. Data represents n = 56, 32, 25, and 35 biologically independent cells respectively, examined from N = 3 independent experimental days. **(F)** Zoom-ins of FAs in the outer region of the cell, as indicated by the boxed regions in (C). Scale bar, 5 µm. To independently examine the effect of the mutations on focal adhesion recruitment, we analysed localisation of VD1 and full-length vinculin mutants in mouse embryonic fibroblasts (MEFs) (**Fig. S8**). Expression of A50I VD1 reduced enrichment in focal adhesions compared with WT VD1 (p < 0.005), whereas I12K/A50I VD1 showed minimal detectable accumulation at adhesions. Quantification of adhesion-to-cytoplasm fluorescence intensity ratios confirmed a strong reduction in VD1 recruitment (WT = 5.14, A50I = 2.08, I12K/A50I = 1.40). In the context of full-length vinculin, both A50I and I12K/A50I reduced focal adhesion enrichment compared with WT vinculin. However, the additional I12K mutation produced only a modest further decrease, with residual adhesion localisation still detectable for the double mutant (adhesion/cytoplasm ratio 1.55 compared with 4.7 for WT). These findings suggest that interactions in addition to talin binding contribute to recruitment and mechanical engagement of full-length vinculin within focal adhesions.

**Figure S8.**
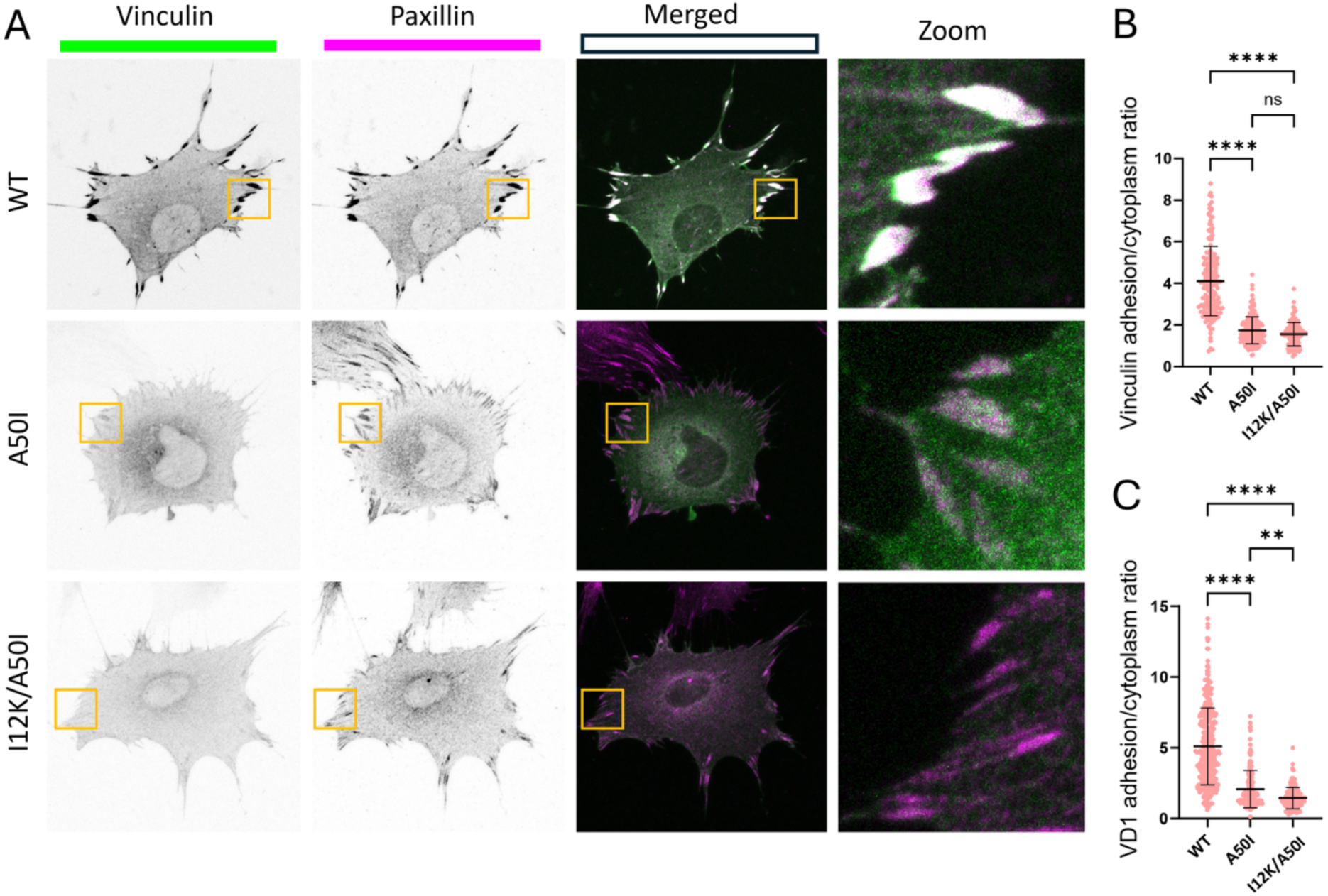
Mutations in VD1 reduce vinculin recruitment to focal adhesions. **(A)** Representative images of MEFs transiently expressing vinculin–EGFP constructs. Paxillin immunofluorescence was used to identify focal adhesions. WT vinculin shows strong co-localisation with paxillin-rich adhesions, whereas A50I vinculin shows reduced accumulation. The I12K/A50I double mutant shows minimal enrichment at adhesions. We note that endogenous vinculin is present in these experiments and the low levels of vinculin mutants at FAs is likely to be enhanced by competition with endogenous vinculin. **(B)** Quantification of vinculin enrichment at adhesions, expressed as the ratio of adhesion to cytoplasmic fluorescence intensity. Approximately 150–200 adhesions from 17–20 cells were analysed across two independent experiments. **(C)** Recruitment of isolated VD1 constructs to focal adhesions. Statistical analysis was performed using ordinary one-way ANOVA. ns, not significant; **p < 0.01; ****p < 0.0001.

### Supplementary Methods relating to Figure S8

#### Cells culture

Wild-type mouse embryonic fibroblasts (MEFs) were maintained at 37 °C in a humidified incubator (85–95 % humidity) with 5% CO₂. The MEF cell line was a kind gift from Dr. Wolfgang Ziegler and has been previously described (Xu et al. 1998). Cells were cultured under sterile conditions in Dulbeccós modified eagle medium (DMEM, high glucose, GlutaMAX™ Supplement, Thermo Fisher Scientific, 61965026,) supplemented with 10% FBS (FBS; Thermo Fisher Scientific, A5256701).

#### Plasmids and mutagenesis

pEGFP-VD1.WT, pEGFP-VD1.A50I, pEGFP-VD1.I12K/A50I, pEGFP-vinculin.FL.WT, pEGFP- vinculin.FL.A50I and pEGFP-vinculin.FL.I12K/A50I plasmids were obtained from GenScript and verified by DNA sequencing. Vinculin inserts were cloned into the pEGFP vector backbone (Clontech).

#### Transfection

Cells maintained in T75 flasks were detached using TrypLE (Thermo Fisher Scientific, 12604013) and centrifuged at 150 g for 5 min. Cell pellets were resuspended in Neon Transfection System R buffer and mixed with 3-6 µg plasmid DNA per reaction.

Cells were transfected by electroporation using a Neon NxT electroporation system (Thermo Fisher Scientific, NEON1S) with the following settings: 1358 V, 30 ms pulse width, 1 pulse. Following electroporation, cells were seeded onto fibronectin-coated coverslips (10 µg/mL, human fibronectin) and allowed to recover and adhere. For time-course experiments, cells were incubated for 8 h or 18 h post-transfection; otherwise, cells were incubated overnight before analysis.

#### Immunofluorescence

To visualise paxillin, transfected cells cultured on coverslips were washed with PBS and fixed in 4% paraformaldehyde (Thermo Fisher Scientific, J61899.AP) for 25 min. Cells were then permeabilised with 0.2% Triton X-100 in PBS for 10 min and washed three times with PBS.

Antibody solutions were prepared in 3% bovine serum albumin (BSA; Sigma-Aldrich, A7906-100G). Cells were incubated with primary antibodies (Table 2) for 45 min using the parafilm method, washed three times with PBS, and then incubated with secondary antibodies for 45 min. Following three additional PBS washes, coverslips were rinsed in Milli-Q water and mounted using ProLong Diamond Antifade Mountant (Thermo Fisher Scientific, P36965).

**Table 2:** Antibodies used for paxillin immunostaining.

| Primary antibody | Dilution | Secondary antibody | Dilution |
| --- | --- | --- | --- |
| Mouse anti-paxillin monoclonal antibody (BD Biosciences, 610051) | 1:100 | Goat anti-mouse Alexa Fluor 568 (Thermo Fisher Scientific, A11004) | 1:250 |

#### Confocal imaging

Imaging was performed using a Zeiss LSM 800 laser-scanning confocal microscope equipped with a 63× oil-immersion objective (Plan-Apochromat, NA 1.40). EGFP-tagged vinculin constructs were excited at 488 nm and Alexa Fluor 568 at 561 nm. Z-stack images were acquired with a 0.23 µm step size between focal planes. Images were collected at 101.4 × 101.4 µm with 8-bit depth.

#### Ratiometric image analysis

To quantify vinculin enrichment at focal adhesions, adhesion-to-cytoplasm EGFP fluorescence intensity ratios were measured using Fiji ImageJ (version 1.54p). Paxillin was used to identify focal adhesions. Regions of interest (ROIs) corresponding to paxillin-rich adhesions were selected using the oval selection tool (8 px brush size), while nearby cytoplasmic regions were selected using a 10 px brush size. Mean fluorescence intensities were measured for each ROI and adhesion-to-cytoplasm ratios calculated. Statistical analyses and graph generation were performed using GraphPad Prism (version 10.3.1).

## Notes

### Competing Interest Statement

The authors have declared no competing interest.

